# Dynamics of fitness distributions in the presence of a phenotypic optimum: an integro-differential approach

**DOI:** 10.1101/444836

**Authors:** M.-E. Gil, F. Hamel, G. Martin, L. Roques

**Affiliations:** Aix Marseille Univ, CNRS, Centrale Marseille, I2M, Marseille, France; BioSP, INRA, 84914, Avignon, France; ISEM (UMR 5554), CNRS, 34095, Montpellier, France

## Abstract

We propose an integro-differential description of the dynamics of the fitness distribution in an asexual population under mutation and selection, in the presence of a phenotype optimum. Due to the presence of this optimum, the distribution of mutation effects on fitness depends on the parent’s fitness, leading to a non-standard equation with “context-dependent" mutation kernels.

Under general assumptions on the mutation kernels, which encompass the standard *n* dimensional Gaussian Fisher’s geometrical model (FGM), we prove that the equation admits a unique time-global solution. Furthermore, we derive a nonlocal nonlinear transport equation satisfied by the cumulant generating function of the fitness distribution. As this equation is the same as the equation derived by Martin and Roques (2016) while studying stochastic Wright-Fisher-type models, this shows that the solution of the main integro-differential equation can be interpreted as the expected distribution of fitness corresponding to this type of microscopic models, in a deterministic limit. Additionally, we give simple sufficient conditions for the existence/non-existence of a concentration phenomenon at the optimal fitness value, i.e, of a Dirac mass at the optimum in the stationary fitness distribution. We show how it determines a phase transition, as mutation rates increase, in the value of the equilibrium mean fitness at mutation-selection balance. In the particular case of the FGM, consistently with previous studies based on other formalisms (Waxman and Peck, 1998, 2006), the condition for the existence of the concentration phenomenon simply requires that the dimension *n* of the phenotype space be larger than or equal to 3 and the mutation rate *U* be smaller than some explicit threshold.

The accuracy of these deterministic approximations are further checked by stochastic individual-based simulations.

## 1 Introduction

Understanding the complex interplay between mutation and selection in asexuals is a central issue of evolutionary biology. Recently, several modeling approaches have been proposed, to describe the evolution of a population under the effects of these two forces. Most of these modeling approaches assume that the fitness of the individuals depends on a single quantitative trait (e.g., [2, 23, 36]). Other approaches, as in this study, directly focus on the dynamics of fitness distributions (e.g., [1, 29]).

A rich literature (reviewed in [33]) has also focused on the interplay between drift selection and mutation in asexuals. These theories focus mainly on the expected mean fitness of the population as it reaches a stationary regime, using models that directly focus on fitness distributions. A few also considered transient behaviors of these models (e.g. [31, 35]), which can be seen as an approximation for the expected distribution of fitness, over time (discussed in [29]).

As fitness is a concept of fundamental importance to our study, we begin with an intuitive definition of this concept. In an asexual population made of *K* genotypes, we say that the genotype *i* has *absolute* Malthusian fitness *m_i_* if the abundance *N_i_*(*t*) of the genotype at time *t* satisfies 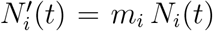. Summing over all the indexes *i* = 1*, …, K*, we observe that the total population size satisfies *N′*(*t*) = *m̄*(*t*) *N* (*t*), with *m̄* (*t*) the mean fitness in the population at time *t*. If we focus on the frequency *p_i_*(*t*) = *N_i_*(*t*)*/N* (*t*), we get 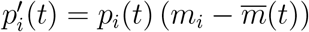.

In this study, following the standard framework of Wright-Fisher or Moran models [25], we assume a constant population size *N* and a continuum of fitness classes *m* ∈ R. In this case, only *relative* Malthusian fitness matters. If we first neglect the effects of mutations, the changes in genotype frequencies due to selection are determined by their relative fitness *m* through the expression (see, e.g., [35]):

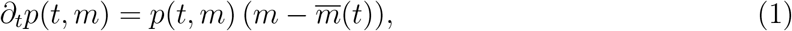

where *m* ∈ ℝ is the relative Malthusian fitness, *p*(*t, m*) the distribution of fitness at time *t*, and *m̄*: ℝ_+_ *→* ℝ is the mean fitness, defined for any *t* ∈ ℝ_+_ by

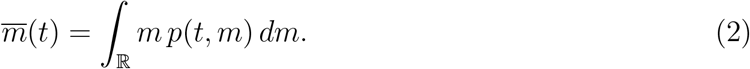

As we can observe in (1), contrarily to other traits, fitness determines its own evolution. We can also note that the notion of relative fitness is defined by (1) up to an additive constant.

A second key step is to describe the distribution of mutation effects on fitness (or DFE, in short). Most mathematical models of asexual evolution (e.g. all those reviewed in [33]) neglect the dependence of the DFE on the fitness of the parent, as discussed e.g. in [18]. Some also ignore deleterious mutations and focus only on the contribution from beneficial ones [33]. However, accumulating data from evolution experiments, often in asexual microbes, has shown that fitness epistasis is pervasive, i.e. that the fitness effects of mutations (individual effects or full distributions) depend on the genetic background in which they arise (e.g. [9, 18, 24]). This has lead to the idea that epistasis, once taken into account, might actually make evolution predictable in spite of stochastic effects due to drift and mutation, at least in large populations and at the fitness level [18, 27, 29]. This may explain why observed fitness trajectories, in large asexual populations, appear relatively repeatable (across biological replicates), in spite of non-repeatable patterns at the sequence level (e.g. [24]). This conjecture partly motivates the present study of phenotypic adaptation as a deterministic dynamical system.

One option to implement epistasis into fitness dynamics models is by using Fisher’s Geometrical Model (FGM) with a single optimum, where a complex form of epistasis arises naturally. The FGM has been shown to lead to fairly realistic DFEs (e.g., [28]) in their shape, dependence on the environment or epistatic pattern (reviewed in [34]). It is a phenotype-fitness landscape model: it assumes *n*-dimensional (breeding values for) phenotype, described by vectors **z** ∈ ℝ^*n*^. The connection between phenotypes and relative Malthusian fitness is made through a quadratic function *m* = −‖**z**‖^2^/2, with ‖ · ‖ the Euclidian norm in ℝ^*n*^. A standard way of describing the effects of mutations on phenotypes is to assume that, given a parent with phenotype **z**, the mutant offspring has a phenotype **z** + *d***z**, where *d***z** follows an *n* dimensional isotropic Gaussian distribution with variance *λ >* 0 at each trait (Gaussian FGM). In this case, even if the distribution of mutation effects on phenotype is independent of the parent phenotype, the DFE on fitness does depends on the parent fitness (because of the non-linear relationship between **z** and *m*). This is illustrated in Fig. 1. These effects can be summarized by a mutation kernel *J_y_*: given a parent with fitness *y*, the mutant offsprings have fitness *y* + *s*, with *s* a random variable with density *J_y_*. We say that this DFE is *context-dependent* because of its dependence on *y*. In spite of this complication, fitness still entirely determines its own evolution, in the FGM under selection and mutation, because epistasis is mediated by fitness alone.

**Figure 1:**
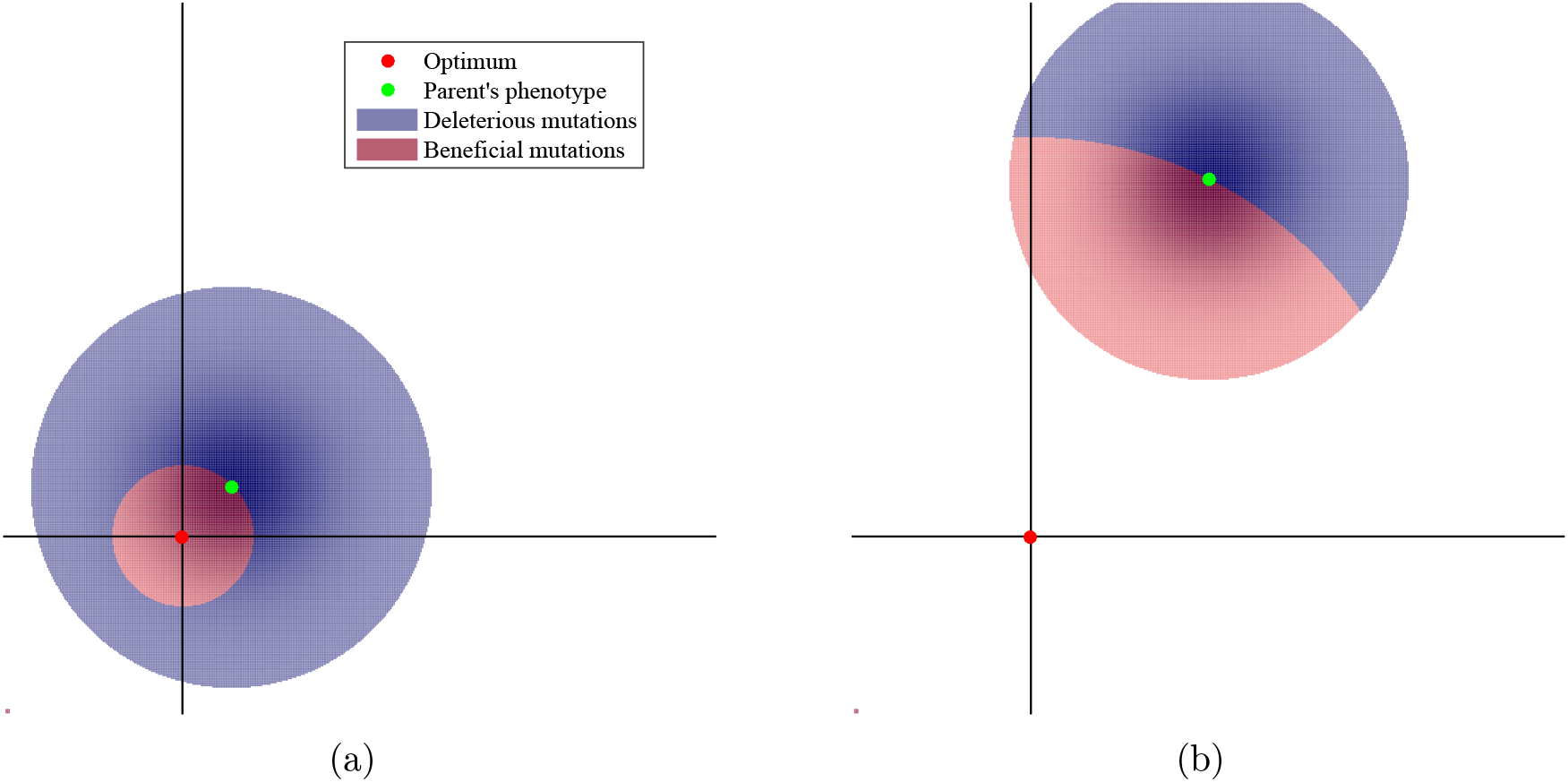
Proportions of beneficial and deleterious mutations, in a two-dimensional phenotypic space with a phenotypic optimum. In panel (b) the parent’s phenotype is farther from the optimum than in panel (a), so that beneficial mutations are more frequent and of larger effect on average in (b).

Combining the equation (1) with the general assumption of a DFE that depends on background fitness, we obtain the following integro-differential equation:

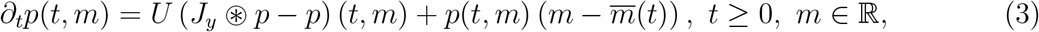

where *U >* 0 is a given constant corresponding to the mutation rate and *J_y_* ⊛ *p* is defined, for any (*t, m*) ∈ ℝ_+_ *×* ℝ, by

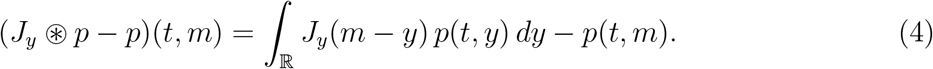

This equation corresponds to a generalization of the equations studied in [17], where *J_y_* = *J* was supposed to be independent of the fitness *y* of the parent, leading to a standard convolution product instead of the operator ⊛. These previous approaches with context-independent DFEs ignore all forms of epistasis, including, for example, that generated by the presence of a fitness optimum. Similarly, other approaches where the mutation effects are modeled with a diffusion approximation, i.e., when *U* (*J_y_* ⊛ *p − p*) (*t, m*) is replaced by *D ∂_mm_p*(*t, m*) for some *D >* 0 also ignore epistasis ([1, 35]). In these two cases (as soon as the support of *J* intersects R_+_, i.e., in the presence of beneficial mutations), the mean fitness *m̄*(*t*) converges to + ∞ at large times, in a non-realistic (superlinear) way, see [31] for a discussion on this aspect.

A closely related work has been proposed in [29]. Their study focuses on a stochastic individual-based Wright-Fisher model combined with the FGM for the description of mutation effects on fitness. Based on formal computations, they derived nonlocal nonlinear transport equations satisfied by some generating function of the fitness distribution. Under some assumptions corresponding to a diffusion approximation of the mutation effects on phenotype, approximate linear transport equations arise that can be solved analytically, allowing to infer the corresponding dynamics of the multivariate phenotype and fitness distribution. Another related work has been developed by [2]. Their approach describes the dynamics of the distribution of a 1-dimensional trait *x*, with corresponding fitness value −*x*^2^, i.e., in the presence of a fitness optimum at *x* = 0. They managed, under a diffusion approximation for the mutation effects on phenotypes, to give a full analytical description of the dynamics of the trait distribution.

From a mathematical perspective, the equation (3) combines several difficulties, compared to standard reaction-diffusion equations *∂_t_u* = *D ∂_xx_u* + *f* (*u*) with local diffusion and local reaction terms. The mutation term *U* (*J_y_* ⊛ *p−p*) is nonlocal, has no regularizing properties, and is not a standard convolution product. The selection term (1) is also nonlocal due to the term *m̄*(*t*). Equations of the type *∂_t_u* = (*J* ⋆ *u u*) + *f* (*u*), with ⋆ the convolution product and a local reaction term *f* (*u*), have been extensively studied, especially regarding the existence/nonexistence of traveling wave solutions and other spreading properties [3, 7, 11, 14, 15, 32, 39, 40, 41]. In the work [17] that was mentioned above, we considered nonlocal reaction terms of the form (1), but again with a standard convolution product. In the recent work [5], a reaction-diffusion equation with a general reaction term of the form 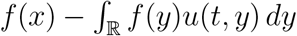 with *f* (*x*) → −∞ as |*x|*→∞, generalizing the results in [2], has also been thoroughly studied. Reaction-diffusion equations with other types of nonlocal reaction terms have also been investigated in recent works [4, 6, 12, 13, 16, 19, 21]. Lastly, we mention that operators of the type *J_y_* ⊛ *p* − *p* or more generally of the type 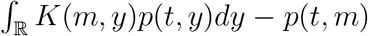 have also been considered in [8, 10] with an emphasis on the study of the stationary states. Moreover, the existence of stationary states and travelling waves for equations of the form *∂_t_u* = *∂_xx_u* + *μ*(*M* ⊛ *u − u*) + *f* (*u*), with *f* (*u*) a nonlocal reaction term, has been studied in [20].

The paper is organized as follows. In Section 2, we detail all the assumptions on the initial condition *p*(0, ·) = *p*_0_ and on the mutation kernels *J_y_*. In Section 3, we present our main results on equation (3). In particular, we derive some a priori estimates on the solutions (Section 3.1); we state an existence and uniqueness result (Section 3.2); we connect this equation with the formal theory developed in [29] for asexual models with selection and epistatic mutation (Section 3.3); we give a qualitative description of the stationary states of (3), and we apply these results to the particular case of the Gaussian FGM (Section 3.4). In Section 4, we present some numerical computations of the solutions of (3) under the assumptions of the Gaussian FGM, and we compare the corresponding distributions with observed distributions from individual-based simulations of the Wright-Fisher model. Lastly, we discuss our results in Section 5. Our results are proved in Section 6.

## 2 Assumptions

### Initial condition

We assume throughout this paper that the initial distribution of fitness *p*_0_ ∈ *L^∞^*(ℝ) ∩ *L*^1^(ℝ) is a probability density function, that is,

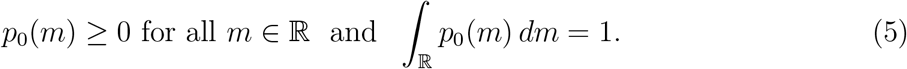

Additionally, we assume that *p*_0_ satisfies

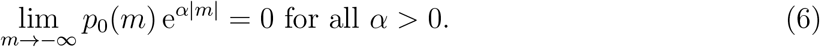

As mentioned in the introduction, we assume that there is a fitness optimum *m*^⋆^ and, without loss of generality as we work with relative fitnesses, we can assume that *m*^⋆^ = 0. Thus, at *t* = 0, all fitnesses must be less than or equal to 0. This means that the initial distribution *p*_0_ is such that:

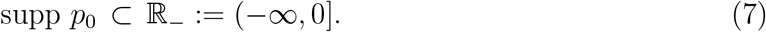

### General mutation kernels

We begin with the general assumptions that we use for our existence and uniqueness result. For each fitness *y* ∈ ℝ_−_, we assume that *J_y_* ∈ *L*^1^(ℝ) is a probability density function:

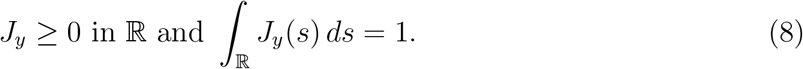

For mathematical convenience, we also set *J_y_* = 0 in R for each *y* ∈ (0, + ∞) and we assume that, for each *m* ∈ ℝ, the function *y* ↦ *J_y_*(*m − y*) is measurable in ℝ and finite almost everywhere (a.e. for short) in R. As 0 is the fitness of the optimum, mutant offspring from any parent with fitness *y* cannot have a fitness larger than 0. It follows that, for each *y* ≤ 0, *−y* is an upper bound of the support of the mutation kernel *J_y_*, that is,

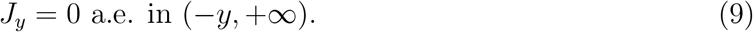

When *y* = 0, the parent has the optimal fitness and all mutations lead to fitnesses *m* ≤ 0. This is consistent with the assumption (9): the kernel *J*_0_ is supported in (−∞, 0], and therefore leads to deleterious mutations only.

For technical reasons, we may also assume that the kernels (*J_y_*)_*y≤*0_ are uniformly bounded in ℝ by a nonnegative function *J̅ ∈ L*^1^(ℝ) which decays faster than any exponential function at −*∞*, in the sense that

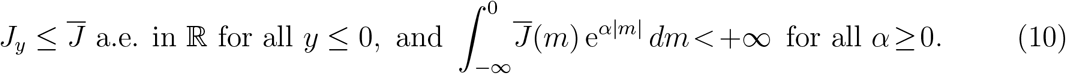

### Kernels with log-linear moment generating function

In order to connect our results with the theory described in [29], and to derive additional properties on the stationary states of (3), we require some additional assumptions on the kernels *J_y_*. We assume here that these kernels have log-linear moment generating function in the sense that

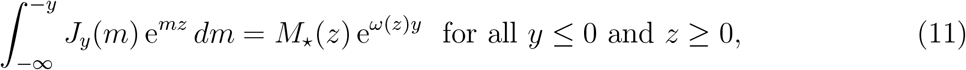

where

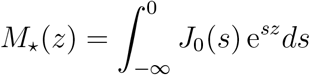

is the moment generating function of the mutation kernel at the optimal fitness and *ω ∈ C*^1^(ℝ_+_) satisfies

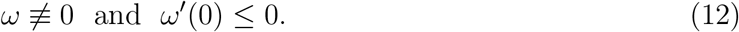

The assumptions (11)-(12) are satisfied in several standard models (discussed in [29]). For example, in the Gaussian Fisher’s Geometric model (FGM) mentioned in Section 1, we have [29]:

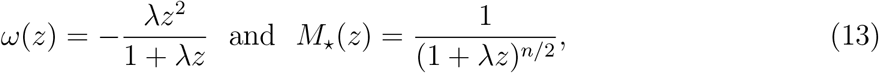

with *λ* > 0 being the variance of mutation effects per phenotypic trait, and *n* the dimension of the phenotype space. The mutation kernels *J_y_* are uniquely defined by their moment generating function *z* ↦ *M*_⋆_(*z*)e^*ω*(*z*)*y*^ and, in the Gaussian FGM, their probability density function (pdf) takes explicit form:

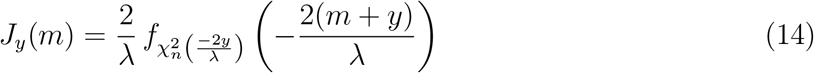

for all *y ≤* 0 and *m ≤ −y*, where 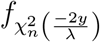 denotes the pdf of the noncentral chi-square distribution with *n* degrees of freedom and noncentrality −2*y/λ*; see Fig. 2 for an illustration. Notice that these kernels also fulfill the assumptions (8)-(10).

**Figure 2:**
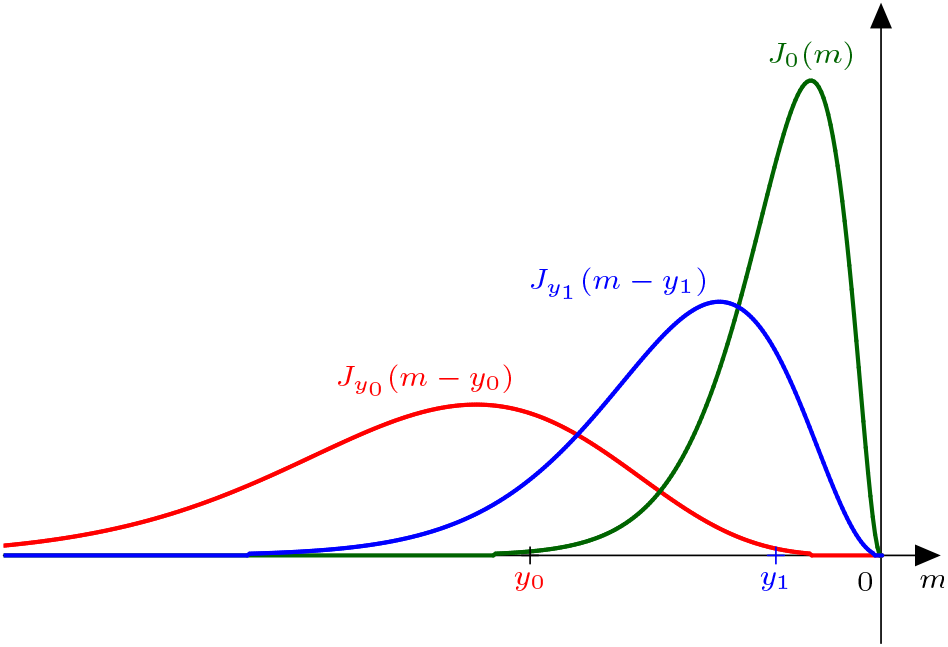
Mutation kernels *J_y_* corresponding to the Gaussian Fisher’s geometric model, with phenotype dimension *n* = 6 and variance *λ* = 1, for parents with fitnesses *y*_0_ *<* 0, *y*_1_ *<* 0 and 0.

The assumption *ω* ≢ 0 in (12) simply means that the kernels *J_y_* do depend on the fitness of the parents. The assumption *ω′*(0) ≤ 0 in (12) can be interpreted as follows. Using (11), we get that:

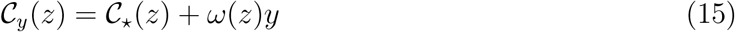

for all *y ≤* 0 and *z ≥* 0, where

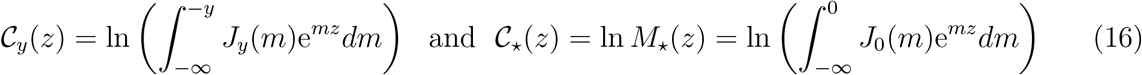

are the cumulant generating functions of *J_y_* and *J*_0_ respectively. Then, from assumption (10), the functions *C_y_* and *C*_*_ = *C*_0_ are of class *C^∞^*(R_+_) and they satisfy

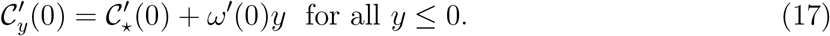

It follows that the mean effect of mutations on fitness *y*, namely 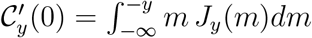, is a non-increasing function of the fitness of the parent 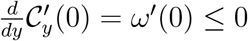. In the special case of the FGM (Fig. 1), the parent fitness does not affect the expected fitness effect of mutations (*ω′*(0) = 0).

## 3 Main results

### 3.1 Support of the solution

Under the assumption (9) on the mutation kernel, it is natural to expect that the upper bound of the support of the solution *p* of (3) remains below the fitness optimum 0. As stated by the proposition below, this is true even without assumptions (6) and (10) on the exponential decay of *p*_0_ and *J_y_* at *−∞*.

#### Proposition 3.1

*Assume that p*_0_ ∈ *L^∞^*(ℝ) ⋂ *L*^1^(ℝ) *satisfies assumption* (7) *and the mutation kernels J_y_*∈ *L*^1^(ℝ) *satisfy assumptions* (8) *and* (9)*. Then, for any T* ∈ (0, +∞] *and any nonnegative solution p of* (3) *such that*

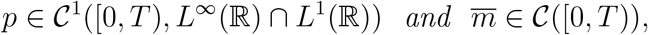

*there holds*

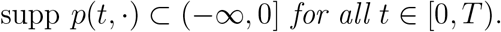

The assumptions and conclusion of Proposition 3.1 show that the integral in (2) is computed over (−∞, 0], as is the integral in (4), since *J_y_* = 0 in ℝ for all *y* > 0. From now on, we may therefore define *m* and *J_y_* ⊛ *p* as

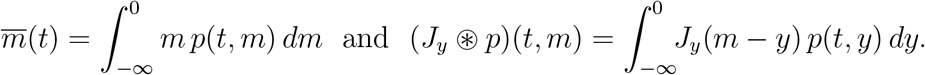

Moreover, for every *t* ∈ [0*, T*), the function (*J_y_* ⊛ *p*)(*t*, ·) belongs to *L*^1^(ℝ) and therefore the integral defining (*J_y_* ⊛ *p*)(*t, m*) converges for almost every *m* ∈ R.

Furthermore, the distribution *p*(*t*, ·) may or not reach the fitness optimum 0. The proposition below shows that if the kernels *J_y_* include some beneficial mutations for any *y <* 0, in some strong sense, then the optimum is instantaneously reached. In other words, the following result gives sufficient conditions for the upper bound of the support of the solution *p*(*t*, ·) to be equal to 0.

#### Proposition 3.2

*Assume that the mutation kernels J_y_* ∈ *L*^1^(ℝ) *satisfy assumptions* (8)*-*(9) *and that the map*

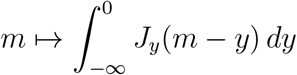

*belongs to L*^∞^(ℝ)*. For any y <* 0*, set*

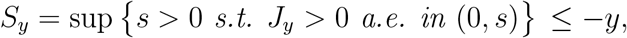

*and assume that*

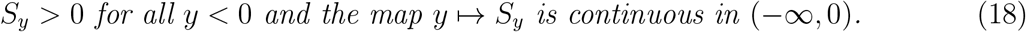

*Assume that p*_0_ ∈ *L*^∞^(ℝ) *∩ L*^1^(ℝ) *satisfies assumptions* (5) *and* (7) *and assume that, for some T* ∈ (0, +*∞*]*, equation* (3) *admits a nonnegative solution p* ∈ *𝒞*^1^([0*, T*), *L^∞^*(ℝ) *∩ L*^1^(ℝ)) *such that m̅* ∈ *𝒞* ([0*, T*)) *and t 1* ⟼ *p̃*(*t, ·*) ∈ *C*^1^([0*, T*), *L^∞^*(ℝ)) *with p̃* (*t, m*) = e^−*tm*^*p*(*t, m*). *Then*

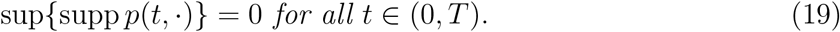

The above result is reminiscent of the strong parabolic maximum principle in parabolic equations (see e.g. [30]): it shows that, provided that each kernel *J_y_* (for *y <* 0) includes some beneficial mutations in the sense of (18), the support of the solution propagates with infinite speed, so that it instantaneously reaches the optimum *m* = 0. This property may not be satisfied without the assumption (18).

### 3.2 Global existence and uniqueness

We are now in position to state our existence and uniqueness result.

#### Theorem 3.3 (Existence, uniqueness, exponential decay)

*Assume that p*_0_ ∈ *L^∞^*(ℝ) ⋂ *L*^1^(ℝ) *satisfies assumptions* (5)*-*(7) *and the kernels J_y_ satisfy assumptions* (8)*-*(10)*. Then problem* (3) *with initial condition p*_0_ *admits a unique solution p ≥* 0 *such that*

*(i)* *t ↦ p*(*t*, ·) ∈ *𝒞*^1^([0, +*∞*), *L^∞^*(ℝ) *∩ L*^1^(ℝ)), *m̄* ∈ *𝒞* ([0, +*∞*)), supp *p*(*t*, ·) ⊂ (*−∞*, 0] *for all t* ∈ [0, +*∞*), *and*

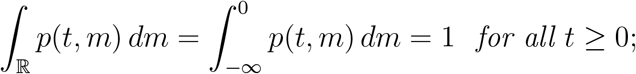
*(ii)* *p decays faster than any exponential function as m*→−∞*in the sense that, for every α >* 0 *and T >* 0*, there is* Γ_*α,T*_ > 0 *such that:*

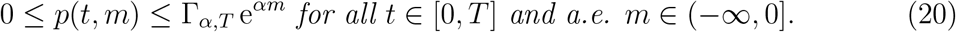 *Lastly, the same decay property* (20) *is valid for |∂_t_p*(*t, m*)*| as well.*

The decay assumptions (6) and (10) on *p*_0_ and *J_y_* seem purely technical in the proof. It is an interesting but still open question to prove the same type of result without these hypotheses.

### 3.3 Cumulant generating function

Our goal here is to connect the equation (3) with the formal theory developed in [29] for Wright-Fisher individual-based models. In [29], the authors derived a nonlocal transport equation approximately satisfied by the expected *cumulant generating function* (CGF, for short) of the fitness distribution.

In the sequel, we assume that the kernels *J_y_* satisfy the assumptions (11)-(12), in addition to the properties (8)-(10). Under the assumptions and notations of Theorem 3.3, we consider the nonnegative solution *p* ∈ *𝒞*^1^([0, +*∞*)*, L^∞^*(ℝ) *∩ L*^1^(ℝ)) of (3), and we define the cumulant generating function *C* ∈ *𝒞*^1^([0, +*∞*) *×* [0, +*∞*)) of the fitness distribution by

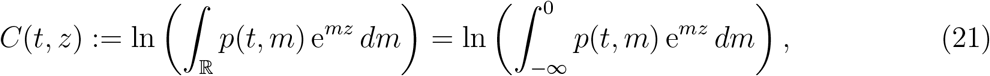

for *t* ≥ 0 and *z* ≥ 0. Notice that, from Theorem 3.3, the map *t* ↦ *𝒮*(*t*,) is actually of class *𝒮*^1^([0, +∞)*, 𝒮^∞^*([0, +∞))). Notice also that the quantity *C* (*t, z*) could be defined for all *t* ≥ 0 and all *z ∈* ℝ due to the decay properties (20). The CGF is a very useful tool to analyze the properties of a distribution. In particular, it is easily seen that

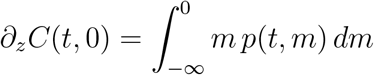

is the mean fitness *m̅*(*t*) at a time *t* ≥ 0 and

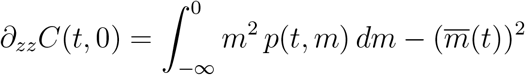

is the variance in fitness within the population.

We now derive the equation satisfied by *C*. For any given *t* ≥ 0 and *z* ≥ 0, by multiplying equation (3) by e^*mz*^ and integrating over (*−∞*, 0] with respect to *m* (all integrals below converge due to the decay properties of *p*(*t, ·*) and *∂_t_p*(*t, ·*) given in Theorem 3.3), we obtain:

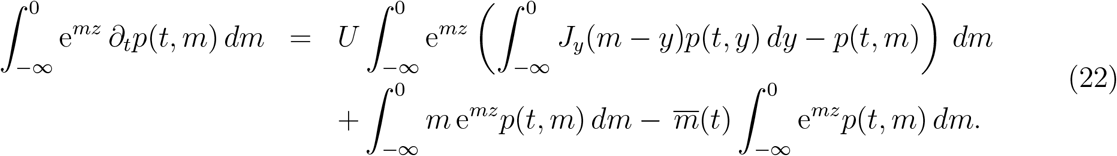

Then, using the assumption (11), we get that:

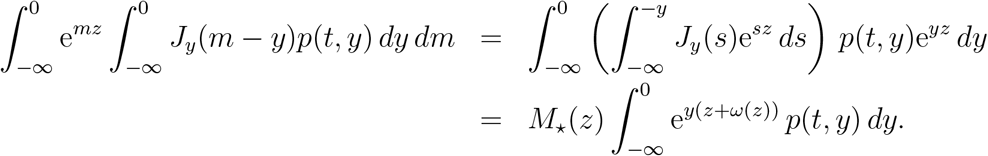

Dividing the equality (22) by 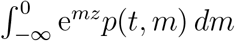 and using Lebesgue’s dominated convergence theorem, we then get the following proposition. In the statement, we point out that *z*+*ω*(*z*) *≥* 0 for all *z ≥* 0, as follows from the assumptions (8), (9) and (11), see (71) and Appendix B below.

#### Proposition 3.4

*The function *C* ∈ 𝒮*^1^([0, +∞)×[0, +∞)) *is a classical solution of the following nonlocal equation*

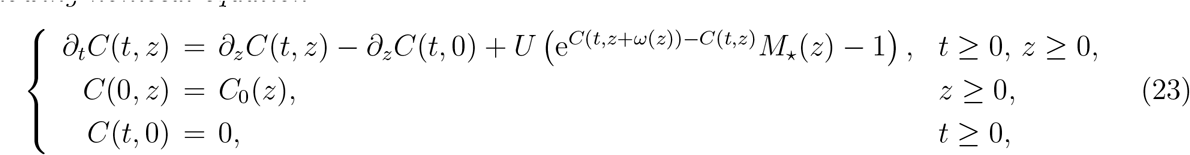
*with initial condition*

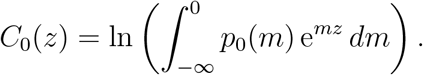

We obtain the same equation as equation (7) in [29]. The consequences of this result are twofold. First, it shows that the solution *p*(*t, m*) of (3) can be interpreted as the expected (expectation among replicate populations) distribution of fitness corresponding to Wright-Fisher-type individual-based models, in a deterministic limit. Second, it gives a rigorous mathematical basis to the statements in [29], which were based on formal computations. In particular, it shows that the solution of (23) (provided that it is unique) is truly a cumulant generating function, in the sense that it satisfies (21), with the following immediate consequences: (i) *C*(*t*, ·) is convex (from Theorem 3.3 and the Cauchy-Schwarz inequality, see (80) below), (ii) *C*(*t*, ·) is a non-increasing function. These two properties were conjectured without proof in [29].

### 3.4 Stationary states

This section is devoted to the study of the stationary states of equation (3). Namely, we focus on weak solutions *p_∞_* of

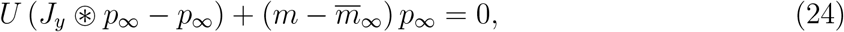

where

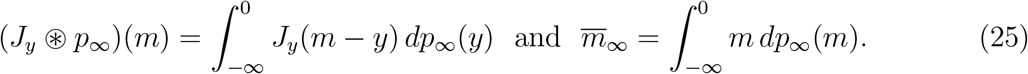

In the section, in addition to the assumptions (5)-(12), we assume that the family (*p*(*t, ·*))_*t≥*0_ converges weakly as *t →* +*∞* to a Radon measure *p_∞_* in the sense that:

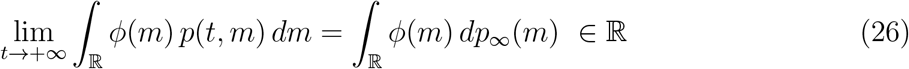

for any continuous function *ϕ*: ℝ *→* ℝ such that *|ϕ*(*m*)*|* = *O*(*|m|*) as *|m| →* +*∞*. It then follows from Proposition 3.1 and Theorem 3.3 that *p_∞_* is nonnegative and satisfies

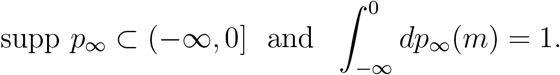

Notice also from (26) that *m̄_∞_* defined in (25) is a real number. Furthermore, the following properties hold.

#### Proposition 3.5

*Assume that* (5)*-*(12), (18) *and* (26) *hold and that, for any continuous function ϕ*: ℝ *→* ℝ *with compact support, the function* (*−∞*, 0] ∋ *y* ⟼ *ψ*(*y*) = ∫_ℝ_ *J_y_*(*m − y*) *ϕ*(*m*) *dm is continuous. Then the measure p_∞_ satisfies*

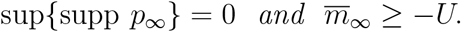

We are now interested in the existence of a positive mass at the optimal fitness (that is, To do so, notice that the measure *p_∞_* can be written as a sum of two measures:

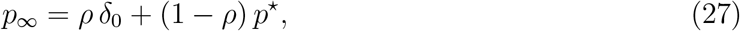

where *ρ* ∈ [0, 1], *δ*_0_ is the Dirac measure at 0, and *p*^⋆^ is a nonnegative measure supported in (*−∞*, 0] such that *p*^⋆^ ((*−∞*, 0]) = 1 and *p*^⋆^ has no mass at 0 in the sense that *p^*^*([*−ε*, 0]) *→* 0 as *ε →* 0. The following result shows a relationship between the existence of a positive mass at 0 and the value of the equilibrium mean fitness *m̄_∞_*.

#### Proposition 3.6

*Under the same assumptions as in Proposition* 3.5*, the measure p_∞_, written as in* (27)*, satisfies*

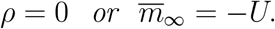

It turns out that the existence of a positive mass at 0 depends on the mutation rate *U* and on the harmonic mean 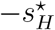 of the kernel at the optimal fitness *J*_0_ defined by

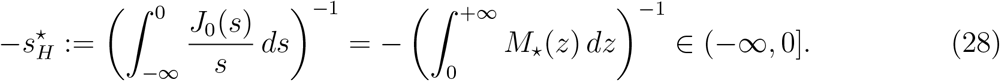

More precisely, the next proposition shows that *p_∞_* admits a positive mass at 0 – meaning that a positive proportion *ρ* of the population has the best possible phenotype – if 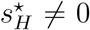 and *U* is not too large.

#### Proposition 3.7

*Under the same assumptions as in Proposition* 3.5*, the measure p_∞_, written as in* (27)*, is such that:*

(i) *if* 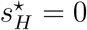, *then ρ = 0; furthermore, if* 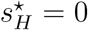 *and*

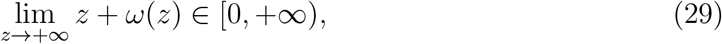

*then m̄_∞_ >* −*U;*
(ii) *if 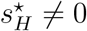, then*
  (a) *if 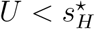, then ρ >* 0 *and m̄_∞_* = *− U;*
  (b) *if*

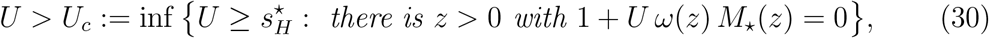

*then ρ* = 0 *and m̄_∞_ > −U.*

These results on the conditions of existence of a positive mass *ρ >* 0 at the optimum are not straightforward to interpret intuitively (see also [37, 38] for a discussion of this issue for the particular case of the FGM). If 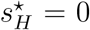 (case (i)), the distribution of mutation effects *s* from the optimal phenotype (kernel *J*_0_) typically decays exponentially or faster around *s* = 0. This means that the optimal phenotype produces an amount of infinitely mild deleterious mutants. These are so mildly counter-selected, relative to their optimal parent, that a population of optimal phenotypes cannot build up, even when the mutation rate is infinitely small (albeit non-zero). On the other hand, when 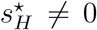 (case (ii)), optimal phenotypes always generate non-vanishingly deleterious mutants, which tend to be counter-selected relative to their optimal parent, thus allowing the maintainance/build-up of a class of optimal phenotypes. However, even then, a large enough mutation rate leads to the erosion of this optimal class (*ρ* = 0), because selection is not strong enough to maintain it, in the face of constant mutation destroying it. This occurs a minima when *U > U_c_* (we point out that *U_c_* is finite from the assumptions made on *J_y_* and *ω*), and cannot occur when 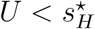.

The assumption (29) on *ω*(*z*) is more technical. As shown in Appendix A, (29) implies that, for every *y <* 0, the upper bound of the support of the kernel *J_y_* is equal to −*y*. In other words, this means that any suboptimal parent (fitness *y <* 0) can yield mutant offspring with the optimal phenotype (fitness 0, mutation effect *s* = −*y*).

Consider finally Fisher’s geometrical model. Namely, assume that the mutation kernels satisfy assumptions (11) and (13)-(14). Then

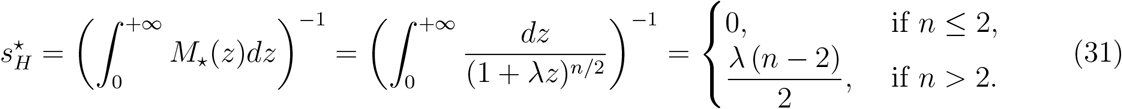

In other words, 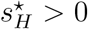 if and only if the phenotype space has more than two dimensions. The following corollary is an immediate consequence of Proposition 3.7 under the assumptions of the Fisher’s geometrical model.

#### Corollary 3.8

*Assume* (5)*-*(14) *and* (26).

(i) *if n ≤* 2*, then ρ* = 0 *and m̄_∞_ >* −*U*;
(ii) *if n >* 2*, then*

(a) *if* 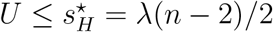, *then, ρ>0 and m̄_∞_ = −U:*
(b) *if* 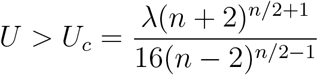, *then, ρ=0 and m̄_∞_ > −U:*

One of the main ingredients in the proofs of the results of this section is the CGF *C_∞_* of *p_∞_* defined by:

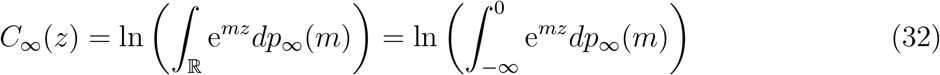

for *z ≥* 0. Assumption (26) shows that *C*(*t, z*) *→ C_∞_*(*z*) as *t →* +*∞* for all *z ≥* 0 and that *C_∞_* is of class *C*^1^([0, +*∞*)) and is convex in [0, +*∞*) as a limit of the convex functions *C*(*t, ·*) in [0, +*∞*). Passing to the limit as *t →* +*∞* in (23), we also obtain that *C_∞_* is a stationary state of (23) in the sense that *C_∞_* is a classical solution of the following equation:

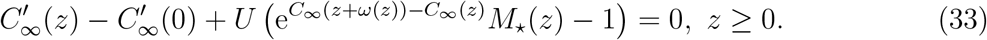

Indeed, for any *s ≥* 0, integrating (23) with respect to *t* over [*s, s* + 1] leads to:

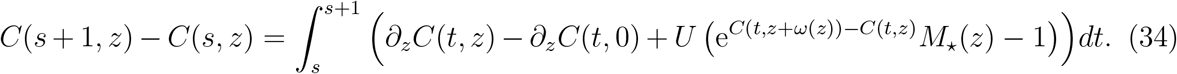

As the function *C* is of class *C*^1^(ℝ_+_ *×* ℝ_+_), the mean value theorem implies that, for each *z ≥* 0 and *s ≥* 0, there is *t_z,s_* ∈ [*s, s* + 1] such that

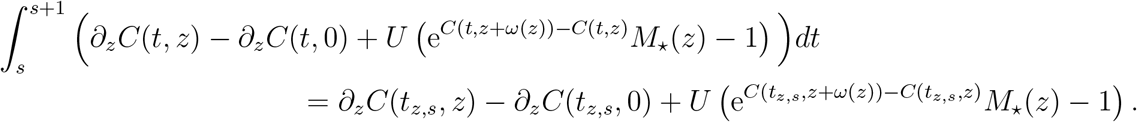

Then, using assumption (26), we get that, for all *z ≥* 0,

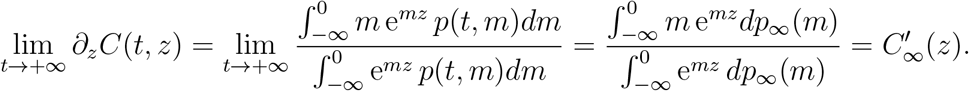

Consequently, passing to the limit as *s* → +∞ in (34) and using lim_*t→*+*∞*_ C(*t, z*) = *C_∞_*(*z*), equation (33) follows. Equation (33) will be used directly in some of the proofs, rather than (24). Notice finally that

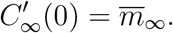

## 4 Numerical computations

The objectives of this section are (i) to check numerically the convergence of the solution of (3) towards an equilibrium; (ii) to illustrate the results of Section 3.4 on the stationary states; and (iii) to compare the distributions *p* obeying the integro-differential equation (3) with empirical individual-based simulations given by a Wright-Fisher model. Namely, we assume that the kernels *J_y_* satisfy the assumptions (8)-(14). For the sake of clarity, we recall these assumptions, which can be summarized as:

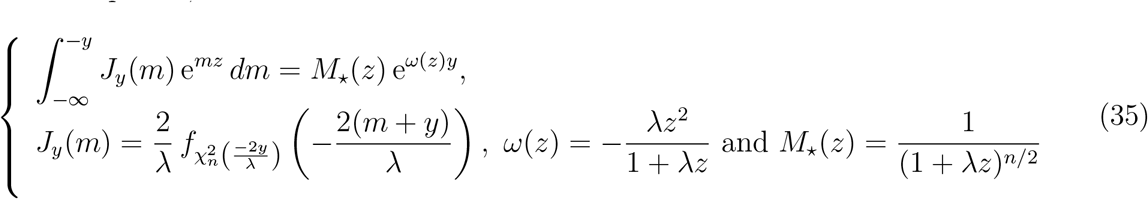

for all *y ≤* 0, *m ≤* −*y* and *z ≥* 0.

### Description of a Wright-Fisher individual-based model

As mentioned in the introduction, we assume a constant population size *N*. Under the assumptions of the FGM, each individual *i* = 1, …, *N* is characterized by a phenotype **z**_*i*_ ∈ ∝ ^*n*^. Its relative Malthusian fitness (exponential growth rate) is *m_i_* = −‖**z**_*i*_‖^2^/2 and its corresponding Darwinian fitness is 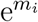 (geometric growth rate, a discrete time counterpart of the Malthusian fitness). We assume non-overlapping generations of duration *δ_t_* = 1. Each generation, selection and genetic drift are jointly simulated by the multinomial sampling of *N* individuals from the previous generation, each with weight given by their Darwinian fitnesses. Mutations are then simulated by randomly drawing, for each individual, a Poisson number of mutations, with rate *U*. We use a classic Gaussian FGM, following e.g. [22, 26]: each single mutation has a random phenotypic effect *d***z** drawn into an isotropic multivariate Gaussian distribution: *d***z ~** *𝒩*(0*, λI_n_*), where *λ >* 0 is the mutational variance at each trait, and *I_n_* is the identity matrix of size *n* × *n*. Multiple mutations in a single individual have additive effects on phenotype. In all our simulations, we started with a clonal population (all of the individuals in the population initially share the same phenotype **z**_0_), and assumed a population size of *N* = 10^5^ individuals.

### Numerical computations

For our simulations, we considered three sets of values of the parameters (*n, λ, U*), each set of value corresponding to a different scenario in Corollary 3.8. The first set of parameters is (*n, λ, U*) = (2, 1/30, 0.05). It corresponds to the assumptions of the first part of Corollary 3.8 (since *n* = 2). The second set of parameters is (*n, λ, U*) = (6, 1/30, 0.05), so that *U < U_c_* ≈ 0.067, corresponding to the second case in Corollary 3.8. The third set of parameters is (*n, λ, U*) = (6, 1/30, 0.55), corresponding to the last case in Corollary 3.8, with *U > U_c_* ≈ 0.533. We solved (3) in a bounded interval [−1.2, 0.05] (but only the values of the solutions in the interval [−0.6, 0.05] are displayed in the figures), using an explicit scheme in time with a time step *δt* = 0.1. The space was discretized, with a uniform step *δm* = 0.001. The simulations were performed using the software Matlab®.

Fig. 3 depicts the dynamics of the fitness distribution obtained with the individual-based and integro-differential approaches, for the three sets of parameter values (*n, λ, U*). In all cases, starting from a clonal population with initial fitness −0.2 (that is, for (3), we consider an initial condition *p*_0_ close to the Dirac mass at −0.2) we observe that the distributions *p*(*t*, ·) obtained by solving (3) seem to converge towards a stationary distribution. The predictions of the integro-differential approach are close to the empirical distribution given by the individual-based model, from qualitative and quantitative viewpoints. Note that, for each set of parameter values, the individual-based simulation was performed only once. The results may vary among replicate simulations.

**Figure 3:**
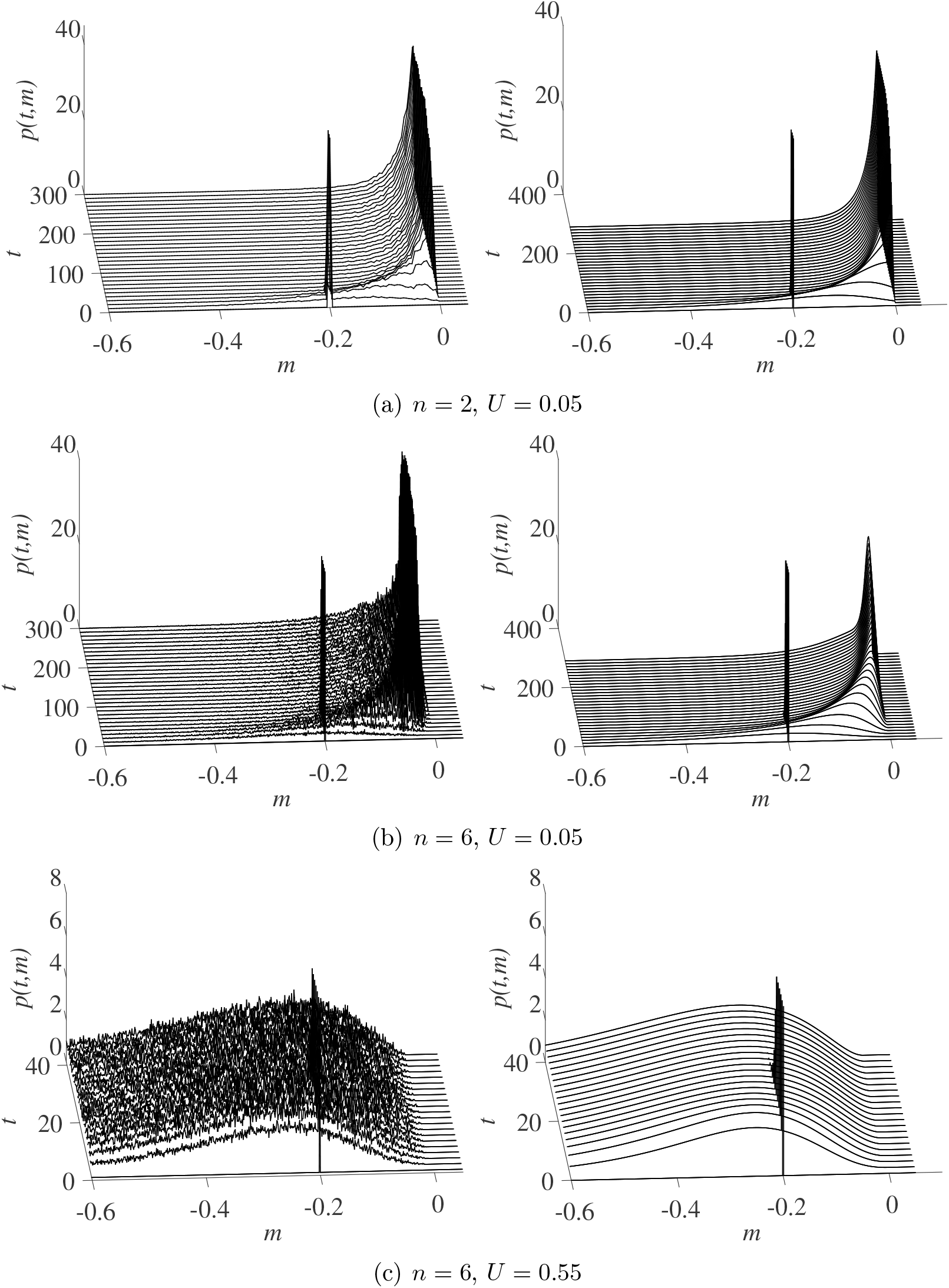
Dynamics of the fitness distribution *p*(*t, m*): individual-based simulation with *N* = 10^5^ individuals (left column) vs numerical solution of (3) with the assumption (35) (right column). In all cases, we fixed *λ* = 1/30 for the mutational variance at each trait, and we assumed a clonal initial population with fitness −0.2.

The predictions of the two approaches at large time (*t* = 5000) are depicted in Fig. 4. Consistently with the result of Corollary 3.8, the distributions in Fig. 4a satisfy *ρ* = 0 and *m̅_∞_ > −U*, the distributions in Fig. 4b satisfy *ρ >* 0 and *m̅_∞_ ≈ −U*, while the distributions in Fig. 4c satisfy *ρ* = 0 and *m̅_∞_ >* −*U*. We also note in all cases a good agreement of the distributions obtained from the integro-differential model with those obtained from the individual-based approach.

**Figure 4:**
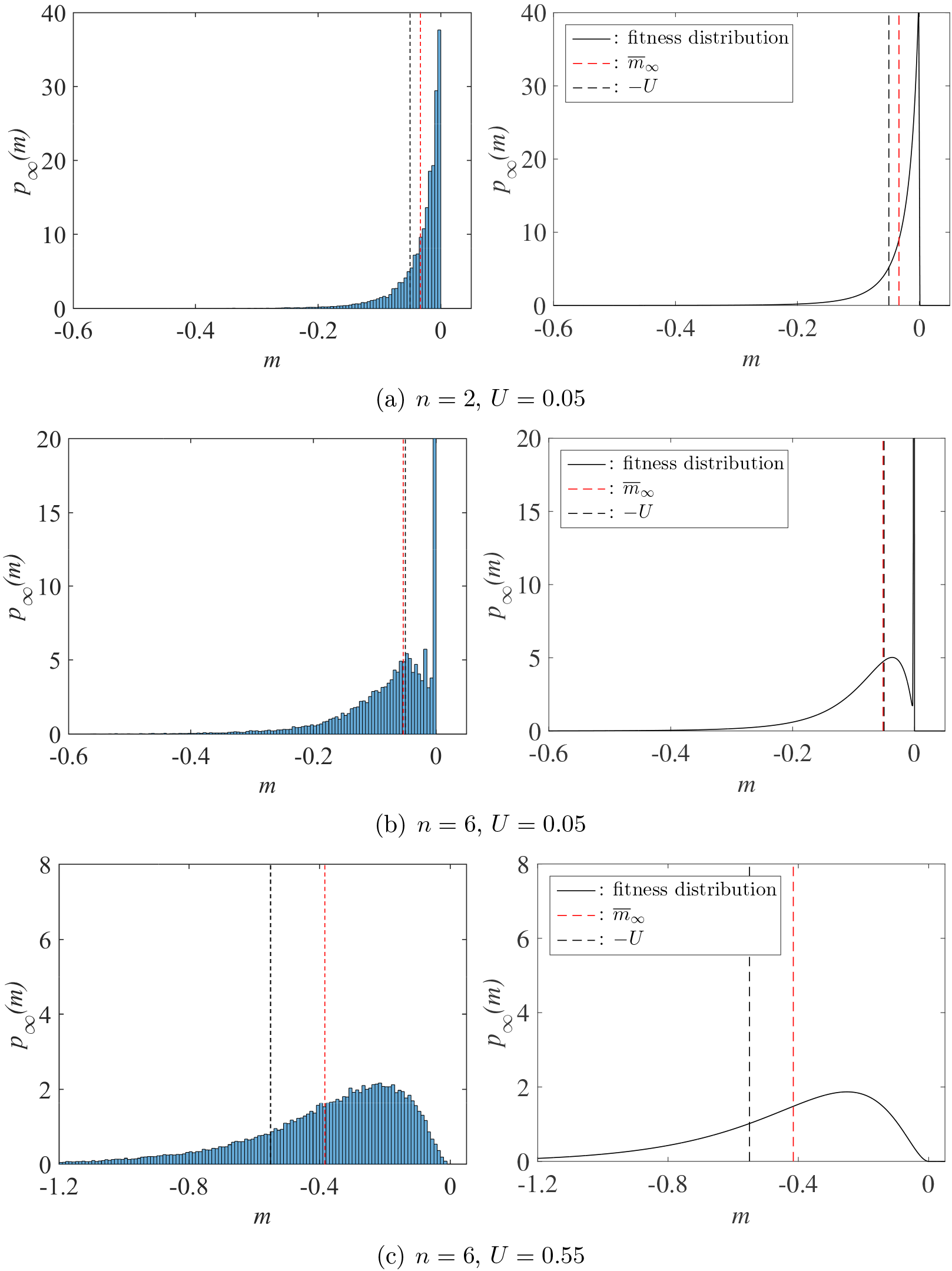
Distribution *p*(*t, m*) at large time *t* = 5000, individual-based simulation (left column) vs numerical solution of (3) with the assumption (35) (right column). The parameter values are the same as in Fig. 3.

## 5 Discussion

We proposed an integro-differential description of the dynamics of the fitness distribution in a population under mutation and selection, in the presence of a phenotype optimum. Under general assumptions on the mutation kernels, which encompass the standard Gaussian Fisher’s Geometrical Model, we proved that the corresponding Cauchy problem (i.e., initial value problem) was "well-posed": it admits a unique time-global solution and the support of the solution remains included in (−∞, 0], consistently with the existence of a fitness optimum at *m* = 0.

Furthermore, we were able to define the cumulant generating function *C*(*t, z*) of the fitness distribution, and to derive a nonlocal nonlinear transport equation satisfied by *C*(*t, z*). This equation is the same as the equation derived in [29] while studying stochastic Wright-Fisher-type individual-based models. We illustrated the connection between equation (3) and a Wright-Fisher-type individual-based model by performing numerical simulations. Under the assumptions of the Gaussian FGM, these simulations showed that equation (3) accurately describes the individual-based dynamics of fitness distributions. Additionally, the simulations suggest that the fitness distribution converges towards a stationary state.

The equation satisfied by *C*(*t, z*) leads to a precise description of these stationary states. In particular, it leads to simple sufficient conditions for the existence/non-existence of a concentration phenomenon at the optimal fitness value *m* = 0 (i.e, of a Dirac mass at *m* = 0 in the stationary fitness distribution). Under the assumptions of the Gaussian FGM, the condition for the existence of the concentration phenomenon simply means that the dimension *n* of the phenotype space is larger than or equal to 3 and the mutation rate *U* is smaller than some explicit threshold. The condition on the dimension of the phenotype space is also a necessary condition: if *n* ≤ 2, a positive mass at 0 can no longer exist. This is reminiscent of the results of [37] who stated that “when three or more characters are affected by each mutation, a single optimal genetic sequence may become common”.

These results on the stationary states also give important clues on the equilibrium value of the mean fitness (*m̄*_∞_) at mutation-selection balance, called "mutation load": if a concentration phenomenon occurs at *m* = 0 then, necessarily, *m̄*_∞_ = −*U*, where *U* denotes the mutation rate. In the absence of concentration phenomenon, we conjecture that *m̄*_∞_ > *U* (this is true in the case of the Gaussian FGM, and in the general case under the technical assumption (29)). The determination of the exact value of *m̄*_∞_ in this case remains a challenging open problem, although approximate treatments are proposed in [29].

The main motivation for this study was to derive rigorous mathematical results when the DFE is given by the Gaussian FGM. It is noteworthy that all of our results remain true for another standard model of context-dependent DFE: the "House of Cards" model. With this approach, given a parent with fitness *y*, the mutant offspring have fitness *s*, where *s* is a random variable with a nonnegative fixed density *J_H_* ∈ *L*^1^(ℝ) supported in (–∞, 0]. This means that the mutation kernels *J_y_* are given by

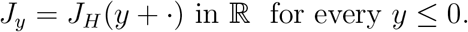

Thus, the family *J_y_* satisfies the same assumption (11) as the Gaussian FGM, with this time

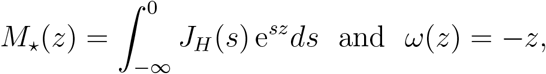

thus, Proposition 3.7 can be applied. In this case, the assumption (29) is always fulfilled, and the occurrence of a concentration phenomenon at the optimum depends on the harmonic mean of *J_H_*.

## 6 Proofs of the main results

This section is devoted to the proof of the main results announced in Section 3 on the solutions of (3), (23) and (33).

### 6.1 Proof of Proposition 3.1. and 3.2 on the support of *p*(*t*, ·)

#### Proof of Proposition 3.1

Assume that *p*_0_ ∈ *L*^∞^(ℝ) ⋂ *L*^1^(ℝ) and the mutation kernels *J_y_* ∈ *L*^1^(ℝ) satisfy assumptions (7)-(9). Let *T* ∈ (0, + ∞] and *p* ∈ *𝒞*^1^([0, *T*), *L^∞^*(ℝ) ⋂ *L*^1^(ℝ)) be a nonnegative solution of (3) such that *m̄* defined by (2) belongs to *𝒞*[0, *T*)). Let *A* > 0 and denote

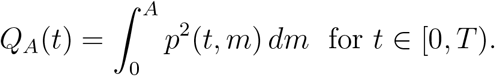

The assumptions on *J_y_* and *p* and Fubini’s theorem imply that, for every *t* ∈ [0, *T*), the function (*J_y_* ⊛ *p*)(*t*,·) belongs to *L*^1^(ℝ) and that

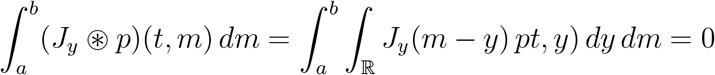

for every 0 *≤ a* < *b*, hence (*J_y_* ⊛ *p*)(*t*, *m*) = 0 for a.e. *m* > 0. Therefore, for every *t* ∉ [0, *T*), by multiplying (3) by *p*(*t*, *m*) and integrating over (0, *A*), we get that

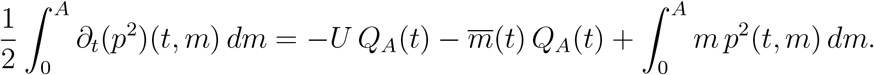

Since

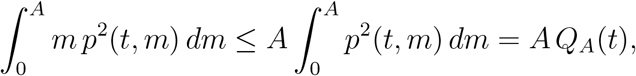

it follows that

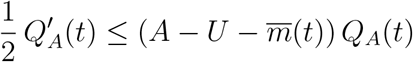

for every *t* ∈ [0, *T*). As *Q_A_*(0) = 0 according to assumption (7), and since *m̄*(*t*) depends continuously on *t* and *Q_A_* ≥ 0 on [0, *T*), Grönwall’s lemma implies that *Q_A_*(*t*) = 0 for all *t* ∈ [0, *T*). Since *A* > 0 can be arbitrarily large, the proof of Proposition 3.1 is thereby complete.

#### Proof of Proposition 3.2

Assume that *p*_0_ and the mutation kernels *J_y_* ∈ *L*^1^(ℝ) satisfy assumptions (5), (7)-(9) and (18). Let *T* ∈ (0, + ∞] and *p* ∈ *𝒞*^1^([0, *T*), *L*^∞^(ℝ) ⋂ *L*^1^(&#z211D;)) be a nonnegative solution of (3) such that *m̄* defined by (2) belongs to *𝒞*([0, *T*)) and *t* ↦ *p̃*(*t*,·) ∈ *𝒞* ^1^([0, *T*), *L^∞^*(ℝ)) with *p̃*(*t*, *m*) = e^*−tm*^p(*t*, *m*). First of all, Proposition 3.1 implies that supp *p*(*t*,·) ⊂ (−∞, 0] for all *t* ∈ [0, *T*).

Define

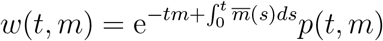

for (*t*, *m*) ∈ [0, *T*) × ℝ. By assumption, the function *t* ↦ *w*(*t*,·) belongs to *𝒞*^1^([0, *T*), *L*^∞^(ℝ)). As *p* obeys (3), *w* is a solution of the following equivalent problem:

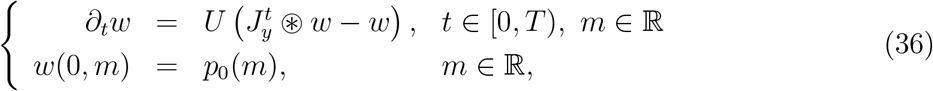

with

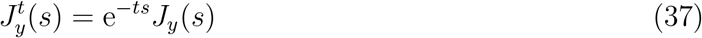

for *t* ∈ [0, *T*), *y* ∈ ℝ and *s* ∈ ℝ. As in (3), the equality 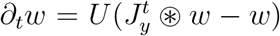 is understood for every *t* ∈ [0, *T*), and for almost every *m* ∈ ℝ (or equivalently for almost every *m* ∈ ℝ_−_ by Proposition 3.1). Observe that, for every *t* ∈ [0, *T*), *p*(*t*,·) and *w*(*t*,·) have the same support (in particular, supp *w*(*t*,) ⊂ (−*∞*, 0] for every *t* ∈ [0, *T*)).

In order to prove that sup{supp *w*(*t*,·)} = 0 for all *t* ∈ (0, *T*), we are going to construct a sub-solution *<u>w</u>* of (36). To do so, assume first, without loss of generality, that *T* < +*∞* (the conclusion in the case *T* = +*∞* would then follow from the case *T* = *n* for any *n* ∈ ℕ, *n* ≥ 1).

Since *p*_0_ is not trivial with *p*_0_ ≥ 0 in R and since supp *p*_0_ ⊂ (−*∞*, 0], there is *μ* < 0 such that

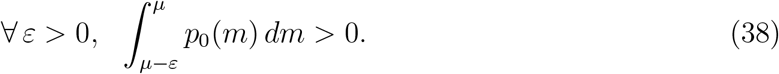

Let now *N* ∈ ℕ such that

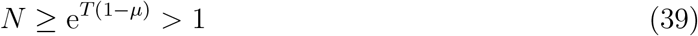

and, for *y* ≤ 0 and *s* ∈ ℝ, define

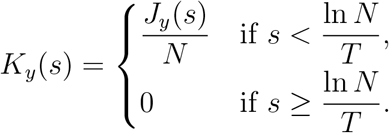

Now, set

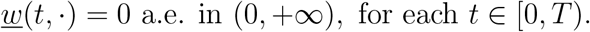

for (*t*, *m*) ∈ [0, *T*) *×* ℝ. Since *p*_0_ ∈ *L^∞^*(ℝ) and the map 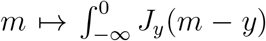 is assumed to be in *L^∞^*(ℝ) as well, one infers that the map *t* ↦ *<u>w</u>*(*t*,·) belongs to *𝒞*^∞^([0, *T*), *L*^∞^(ℝ)). Furthermore, since supp *p*_0_ ⊂ (*−∞*, 0] and *J_y_* = 0 a.e. in (*−y*, +*∞*) for all *y* ≤ 0, one has *K_y_* ⊛ *p*_0_ = 0 a.e. in (0, +*∞*) (as for (*J_y_* ⊛ *p*)(*t*,·) in the proof of Proposition 3.1) and

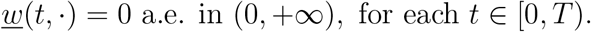

There also holds

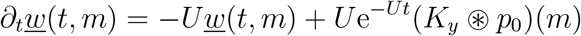

and

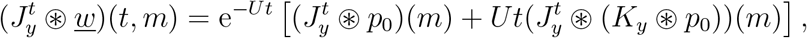

hence

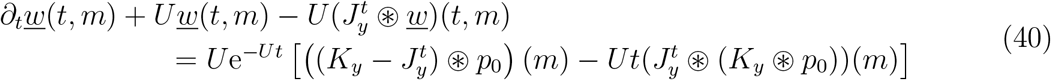

for (*t*, *m*) ∈ [0, *T*) *×* ℝ. Using the nonnegativity assumptions (5) and (8), we get, for each *t* ∈ [0, *T*) and *a < b* in ℝ,

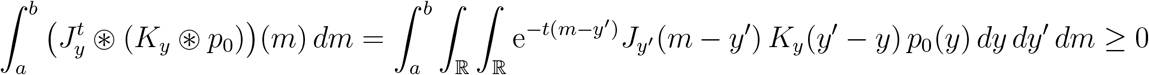

by Fubini’s theorem, hence

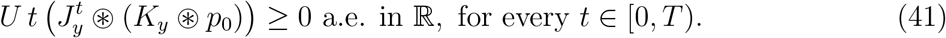

Moreover, for each *t* ∈ [0, *T*), one has 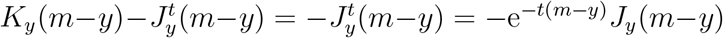 if *m − y* ≥ (ln *N*)*/T*, whereas

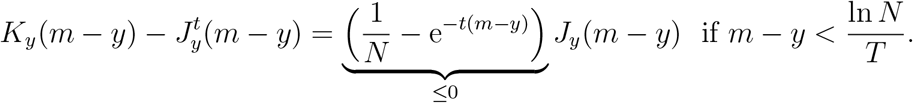

As a consequence, from (8), one infers that, for every *t* ∈ [0, *T*) and *a* < *b* in ℝ,

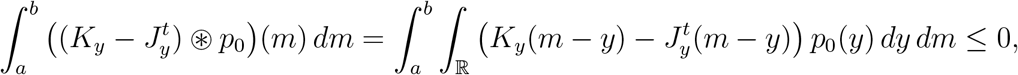

hence 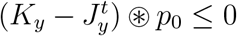 a.e. in ℝ for every *t* ∈ [0*, T*). Then, together with (40) and (41), we get that

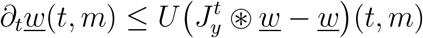

for all *t* ∈ [0, *T*) and for a.e. *m* ∈ ℝ. To sum up, *<u>w</u>* ∈ *𝒞^∞^*([0, *T*], *L^∞^*(ℝ)) is a sub-solution of problem (36).

Remember also that *<u>w</u>*(0,·) = *p*_0_ = *w*(0,·) in ℝ. In order to conclude that *<u>w</u>*(*t*, ·) ≤ *w*(*t*, ·) a.e. in ℝ for every *t* ∈ [0, *T*), we will apply the following comparison principle, whose proof in postponed in Section 7.

##### Lemma 6.1 (Comparison principle)

*Let τ* ∈ (0, +*∞*), *ϖ* ∈ *𝒞* ([0, *τ*]) *and h*_1_, *h*_2_ ∈ *𝒞*^1^([0, *τ*]*, L^∞^*(ℝ)) *be such that, for every t* ∈ [0, *τ*], *h*_1_(*t*,·) = *h*_2_(*t*, ·) = 0 *a.e. in* (0, +*∞*) *and*

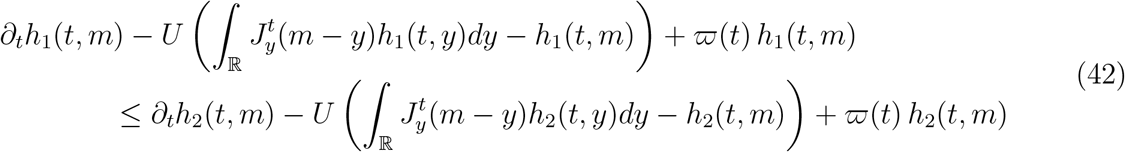

*for a.e. m* ∈ ℝ_−_, *with 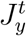 defined in* (37)*. Assume that h*_1_(0,·) ≤ *h*_2_(0, ·) *a.e. in* ℝ*. Then, for every t* ∈ [0, *τ*], *h*_1_(*t*, ·) ≤ *h*_2_(*t*, ·) *a.e. in* ℝ.

Together with the fact that the map *t* ↦ *w*(*t*,·) belongs to *𝒞*^1^([0, *T*), *L*^∞^(ℝ)), Lemma 6.1 (applied with *h*_1_ = *<u>w</u>*, *h*_2_ = *w*, *ϖ* = 0 and every *τ* ∈ (0, *T*)) yields *<u>w</u>*(*t*, ·) ≤ *w*(*t*,·) a.e. in ℝ for every *t* ∈ [0, *T*). Hence, for each *t* ∈ [0, *T*), there holds

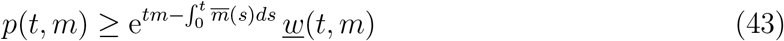

for a.e. *m* ∈ ℝ.

Let now *σ >* 0 be such that

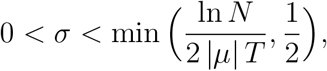

where *μ <* 0 and *N* > 1 were given in (38) and (39). Let us show in this paragraph that, for every *t* ∈ (0, *T*), *<u>w</u>*(*t*,·) > 0 in [*μ*, *μ* + *σS_μ_*], where we recall that *S_μ_ >* 0 from assumption (18). Using assumption (18) on the continuity of (*−∞*, 0) ∋ *y* ↦ *S_y_* ∈ (0, *−y*] and the property (38) on *μ*, there is *β* ∈ (0, *σS_μ_*] such that *S_y_ >* 2 *σ S_μ_* for all *y* ∈ (*μ* − *β*, *μ*) and

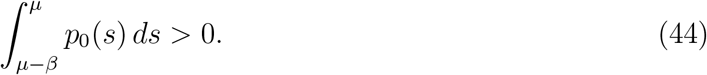

Pick any *t* ∈ (0, *T*) and any *a* < *b* ∈ [*μ*, *μ* + *σS_μ_*]. We have

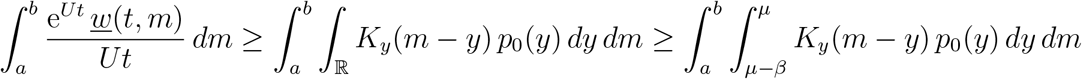

by Fubini’s theorem and the nonnegativity of *K_y_* and *p*_0_. As 0 < *β* ≤ *σS_μ_*, *μ* ≤ *a* < *b* ≤ *μ*+*σS_μ_* and 0 < *S_μ_* ≤ *|μ|*, one infers that, for every *y* ∈ (*μ − β*, *μ*) and *m* ∈ [*a*, *b*],

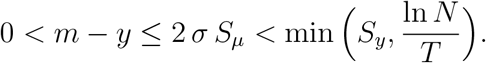

We deduce from the definition of *S_y_* that

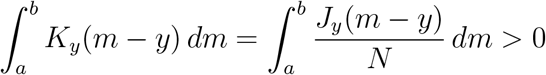

for all *y* ∈ (*μ − β*, *μ*). Together with (44) and the nonnegativity of *p*_0_, we get that

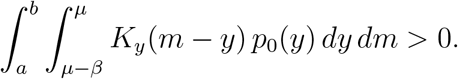

As a consequence,

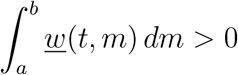

for all *a < b* ∈ [*μ*, *μ* + *σS_μ_*].

Pick any *t*_0_ in (0, *T*) and remember that *μ* < *μ* + *σS_μ_* < 0 since 0 < *σ* < 1/2 < 1 and 0 *< S_μ_* ≤ *−μ* = *|μ|*. It follows from (43) and the previous paragraph that

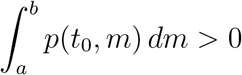
for all *a < b ∈* [*μ*, *μ* + *σS_μ_*] (and *p*(*t*_0_,) ≥ 0 in ℝ by assumption), hence

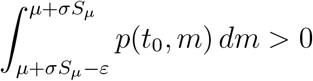

for every *ε* > 0. By observing that 0 < *S_μ+σS_μ__* ≤ −(*μ* + *σS_μ_*) < − *μ* = |*μ|* and by using the same argument as in the previous paragraph with the initial condition *p*(*t*_0_, ·) instead of *p*_0_ and with *μ* + *σS_μ_* instead of *μ*, we get that, for every *t* ∈ (*t*_0_, *T*) and for every 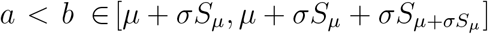,

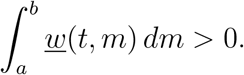

As *t*_0_ is arbitrary in (0, *T*) and by remembering (43), we infer that, for all *t* ∈ (0, *T*) and 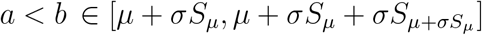,

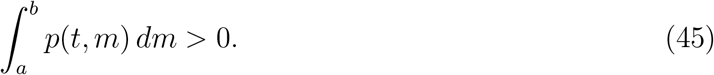

Consider finally the sequence (*S̃_n_*)_*n*∈ℕ_ defined by

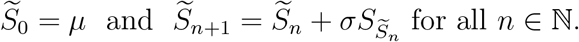

From the inequalities 0 < *S_y_* ≤ −*y* (for all *y* < 0) and 0 < *σ* < 1/2 < 1, the sequence (*S̃_n_*)_*n*∈ℕ_ is well-defined, increasing, and such that *μ* ≤ *S̃_n_ <* 0 for all *n* ∈ ℕ. It then converges to a limit

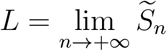

such that *μ* < *L* ≤ 0. If *L* were (strictly) negative, then the continuity assumption (18) of *y* ↦ *S_y_* in (*−∞*, 0) would imply that *L* = *L* + *σS_L_*, whence *S_L_* = 0, which is ruled out by (18). Therefore, *L* = 0. On the other hand, the arguments of the previous paragraphs imply by immediate induction that (45) holds for every *t* ∈ (0, *T*), every *n* ∈ ℕ and every *a < b* ∈ [*μ, S̃_n_*_+1_]. Hence, (45) holds for every *t* ∈ (0, *T*) and *a* < *b* ∈ [*μ*, 0). Together with Proposition 3.1, one gets that

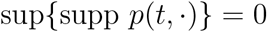

for every *t* ∈ (0, *T*). The proof of Proposition 3.2 is thereby complete.

### 6.2 Global existence: proof of Theorem 3.3

In order to show Theorem 3.3 on the global existence of solutions of (3), the general strategy consists in applying Cauchy-Lipschitz theorem in some suitably chosen function space. To do so, we first prove the local existence, with an existence time which is quantitatively defined in terms of the kernels (*J_y_*)_*y*∈ℝ__ and the initial probability density *p*_0_.

#### Proposition 6.2 (Local existence)

*Let β* ≥ 1, *let the kernels J_y_ satisfy assumptions* (8)*-*(10) *and let p*_0_ ∈ *L^∞^*(ℝ) ⋂*L*^1^(ℝ) *satisfy* (5), (7) *and*

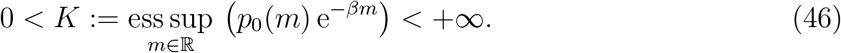

*Let*

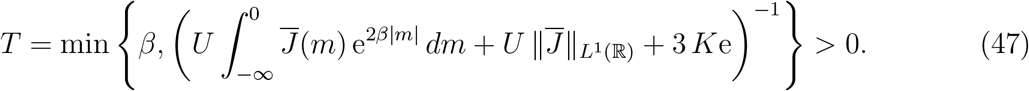

*Then problem* (3) *admits a solution p* ∈ *𝒞*^1^([0, *T*], *L^∞^*(ℝ) ⋂*L*^1^(ℝ)) *such that m̄* ∈ *𝒞*([0, *T*]) *and p decays at least like* e^*t/T* +*βm*+*tm*^ *as m* → −*∞*, *in the sense that*

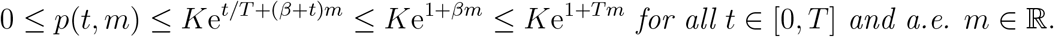

*Furthermore, this solution is unique.*

**Remark 6.3** Notice that supp *p*(*t*, ·) ⊂ ℝ_−_ for every *t* ∈ [0, *T*) from Proposition 3.1 and for *t* = *T* too from the continuity of the map *t* (∈ [0, *T*]) ↦ *p*(*t*, ·) in *L*^∞^(ℝ) ⋂ *L*^1^(ℝ).

#### Proof of Proposition 6.2.

*Step 1: an auxiliary problem*. Let *β*, *p*_0_ and *K* be defined as in the statement of Proposition 6.2. We first show the local existence, for some well chosen *T >* 0, of a solution *υ* of the following nonlinear Cauchy problem:

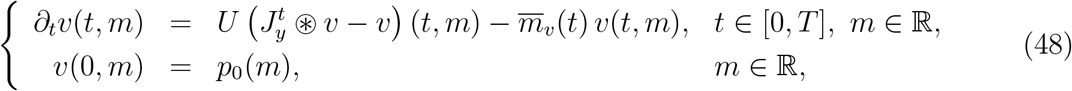

with 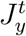 as in (37) and

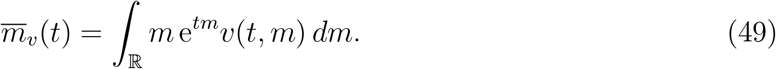

To do so, let us first introduce the Banach space

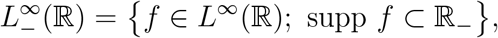

pick any real number *T* such that 0 *< T* ≤ *β*, and consider the set

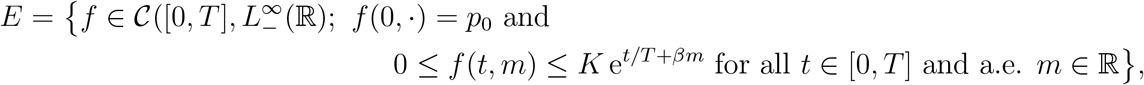

and denote^1^

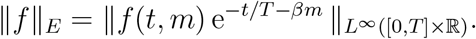

Notice that the set *E* is not empty since the function *f* defined by *f* (*t*, *m*) = *p*_0_(*m*) belongs to *E*, as *p*_0_ satisfies (5), (7) and (46). Let us now define a map *F* as follows:

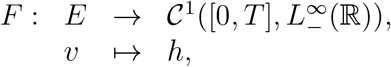

where *h* = *F* (*υ*) is the solution of the following linear Cauchy problem

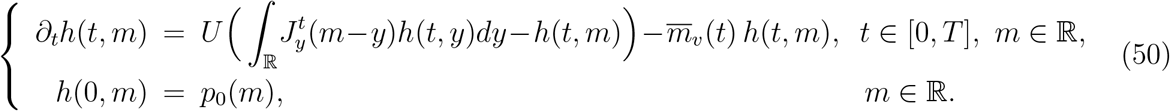

Notice that, since *υ* ∈ *E*, the function *m̄_υ_* defined in (49) exists and belongs to *𝒞*([0, *T*]). Furthermore, Lemma 6.4 below states that *h* is well defined as well.

#### Lemma 6.4

*For any given υ* ∈ *E with* 0 < *T* ≤ *β, the Cauchy problem* (50) *admits a unique solution h* ∈ *𝒞*^1^([0, *T*], *L*^∞^(ℝ)).

In order not to slow down the proof of Proposition 6.2, the proof of Lemma 6.4 is postponed in Section 7.

*Step 2: F maps E to E for T >* 0 *small enough.* Consider any function *υ* ∈ *E*, for some *T* ∈ (0, *β*]. Since supp *υ*(*t*, ·) ⊂ ℝ_−_ for all *t* ∈ [0, *T*] and since 0 ≤ *υ*(*t*, *m*) ≤ *K* e^*t/T* +*βm*^ for all *t* ∈ [0, *T*] and a.e. *m* ∈ ℝ_−_ with *β* ≥ 1, it follows that, for all *t* ∈ [0, *T*],

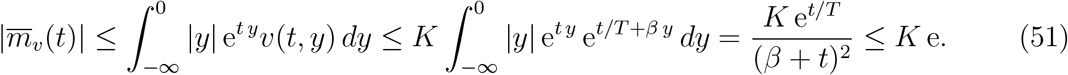

Now, set

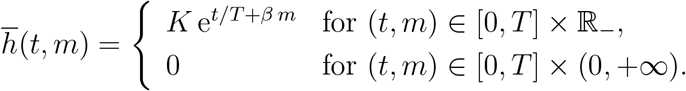

Using assumption (9) and the fact that *J_y_* = 0 in ℝ for all *y* > 0, we observe that

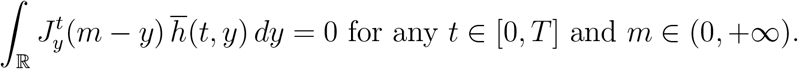

On the other hand, for any (*t*, *m*) ∈ [0, *T*] *×* ℝ_−_, it follows from (10) and (37) that

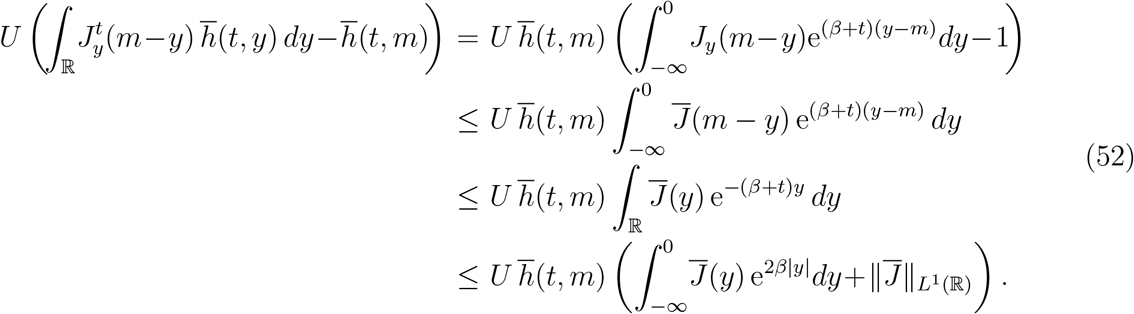

Consider any real number *T* such that

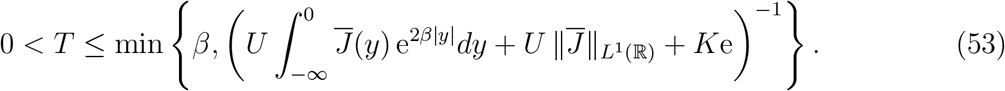

Therefore, using the inequalities of the previous paragraph together with (51), we get that, for all (*t*, *m*) ∈ [0, *T*] *×* ℝ,

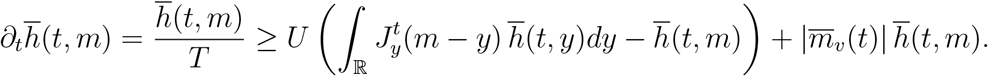

Observe also that 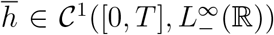. In other words, 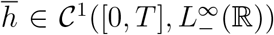 is a supersolution of problem (50) satisfied by *h* = *F*(*υ*). From the definition (46) of *K* and from (7), we also know that 0 ≤ *h*(0, *m*) = *p*_0_(*m*) ≤ *h̄*(0, *m*) for a.e. *m* ∈ ℝ.

Using the comparison principle of Lemma 6.1, we obtain that, for every *t* ∈ [0, *T*], 0 ≤ *h̄*(*t*, *m*) ≤ *h*(*t*, *m*) for a.e. *m* ∈ ℝ. Together with Lemma 6.4, it follows that *h* = *F* (*υ*) ∈ *E*.

*Step 3: F is a contraction mapping*. Let *T >* 0 be as in (53), let *υ*_1_, *υ*_2_ ∈ *E* and define *H* = *F* (*υ*_1_) *− F* (*υ*_2_). Notice that the function *H* belongs to 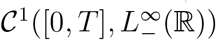 from Lemma 6.4 and that it satisfies, for all *t* ∈ [0, *T*] and a.e. *m* ∈ ℝ,

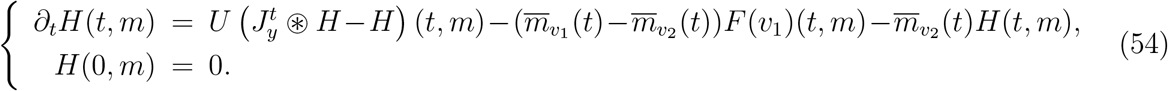

Define

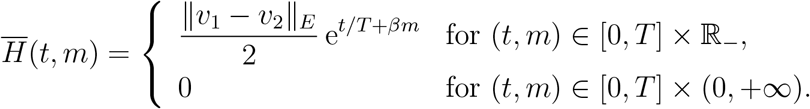

Similar computations as for *h̄* and *υ* above imply that

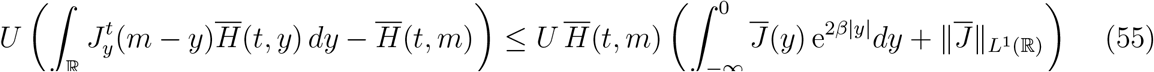

for all (*t*, *m*) ∈ [0, *T*] *×* ℝ, and that 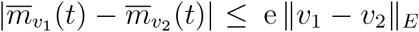 for all *t ∈* [0, *T*]. Since *F* (*υ*_1_) ∈ *E*, it then follows that

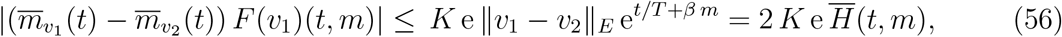

for all *t* ∈ [0*, T*] and a.e. *m* ∈ ℝ_−_.

Finally, let us consider the real number *T* defined by (47). Using (51), (55) and (56), we get that

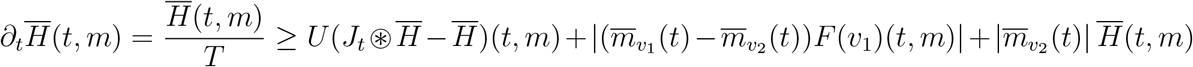

for all *t* ∈ [0*, T*] and *m* ∈ ℝ_−_ (and also for *m >* 0 since all quantities are then equal to 0). As above, this implies that *H̅* is a supersolution of equation (54) satisfied by *H*, while 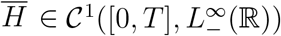. The functions *h*_1_: = *H* and *h*_2_: = *H̅* then satisfy some inequalities of the type (42) with *ϖ* = –*m̅_υ2_* and *τ*: = *T*. Using again the comparison principle of Lemma 6.1, we get that

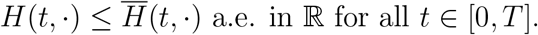

Similarly, one has *−H*(*t, ·*) *≤ H̅* (*t, ·*) a.e. in ℝ for all *t* ∈ [0*, T*]. This immediately yields

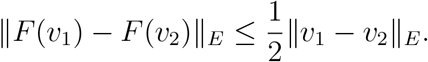

Thus, *F*: *E → E* is a contraction mapping.

*Step 4: conclusion*. Consider now *T >* 0 as defined in (47). The Banach fixed point theorem implies that *F* admits a unique fixed point *υ* ∈ *E*. This function *υ* then belongs to 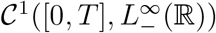 from Lemma 6.4, it satisfies

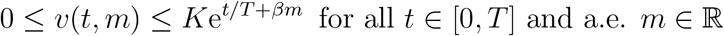

with supp *υ*(*t*, ·) ⊂ ℝ_−_ for every *t* ∈ [0*, T*]. This function *v* is also the unique such solution of (48). Furthermore, with similar estimates as in (51) and (51), there is *K*′ *>* 0 such that |*∂_t_υ*(*t, m*)| ≤ *K*′e^*t/T* +*βm*^ for all *t* ∈ [0*, T*] and a.e. *m* ∈ ℝ_−_, and *∂_t_υ*(*t, m*) = 0 for all *t* ∈ [0*, T*] and a.e. *m >* 0.

Finally, letting

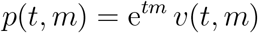

for (*t, m*) ∈ [0*, T*] *×* ℝ, it is straightforward to check that *p* ∈ *𝒞*^1^([0*, T*]*, L^∞^*(ℝ) is a solution of (3) in [0*, T*] *×* ℝ, with supp *p*(*t, ·*) ⊂ ℝ_−_ for every *t* ∈ [0*, T*]. Additionally, as *υ* ∈ *E*, it follows that *p*(*t, ·*) ∈ *L*^1^(ℝ) for all *t* ∈ [0*, T*]. Lastly, *p* ∈ 𝒞^1^([0*, T*]*, L^∞^*(ℝ) *∩ L*^1^(ℝ)), the function

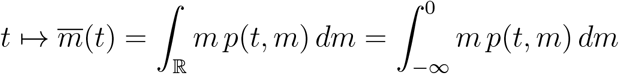

is continuous in [0*, T*]), and

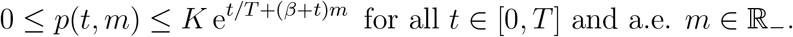

The proof of Proposition 6.2 is thereby complete. D

#### Proof of Theorem 3.3

We are now in position to prove the global existence result of Theorem 3.3. Let *T_max_ >* 0 be the largest time such that equation (3) admits a solution *p* ∈ *𝒞*^1^([0*, T_max_*)*, L^∞^*(ℝ) ∩ *L*^1^(ℝ)) with *m̅* ∈ ([0*, T_max_*)) and, for every *T* ∈ (0*, T_max_*), there is *C_T_ >* 0 such that

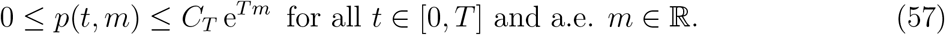

Notice that Proposition 3.1 then implies that supp *p*(*t*, ·) ⊂ ℝ_−_ for every *t* ∈[0*, T_max_*). It follows from the local existence and uniqueness result of Proposition 6.2 applied with, say, *β* = 1 that

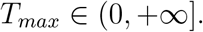

Notice that *T_max_* does not depend on the choice *β* = 1 in Proposition 6.2, since (57) does not involve any *β*. Our goal is to show that *T_max_* = + ∞.

We begin with some fundamental estimates stated in the following lemma, whose proof is postponed in Section 7.

##### Lemma 6.5 (Mass preservation and estimates on the mean fitness)

*We have:*

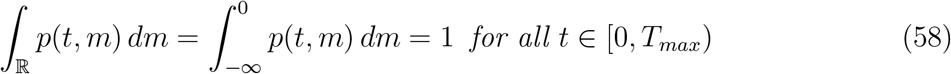

and

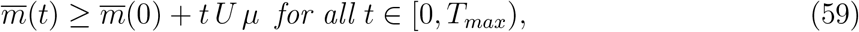

with

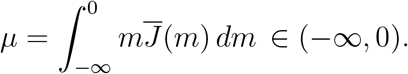

Assume now by contradiction that *T_max_ <* +*∞*. Define

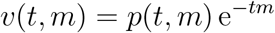

for (*t, m*) ∈ [0*, T_max_*) *×* ℝ. The function *υ* is a solution of the Cauchy problem (48) for all *T* ∈ [0*, T_max_*) and, from (57) and the regularity properties of *p*, the function *υ* 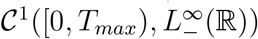. Set now

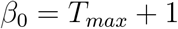

and let

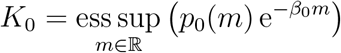

be defined as in (46) in Proposition 6.2 with the choice *β* = *β*_0_. Denote

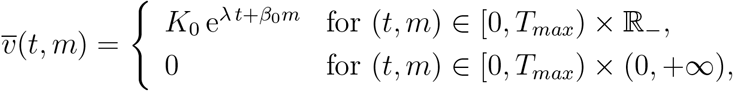

where λ ∈ ℝ is to be chosen later. Using (59) and the property *m̅*(*t*) = *m̅_υ_*(*t*) for all *t* ∈ [0*, T_max_*), it is easily seen (as in Step 2 of the proof of Proposition 6.2) that, for every *T* ∈ (0*, T_max_*), *υ̅* is a supersolution of the equation (48) (for which *m̅_υ_*(*t*) = *m̅* (*t*) is considered as a fixed coefficient) satisfied by *υ* on [0*, T*], provided that

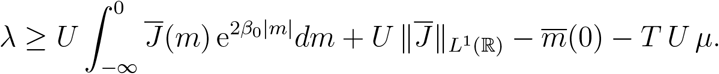

Let then

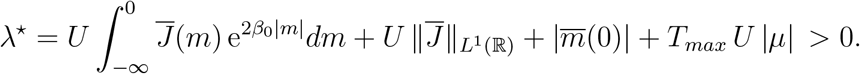

Using the comparison principle of Lemma 6.1 applied with every *τ* ∈ (0*, T_max_*), we obtain that

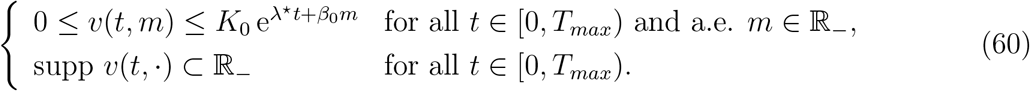

Now, for any *θ* ∈ (0, 1), set *p_θ_* = *p*(*θ T_max_, ·*). We have

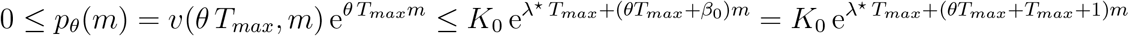

for a.e. *m* ∈ ℝ_−_, while supp *p_θ_* ⊂ ℝ_−_. For any *θ* ∈ (0, 1), the function *p_θ_* satisfies (5), owing to (57) and (58). Furthermore,

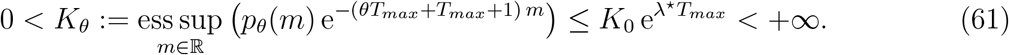

As a consequence, we can apply Proposition 6.2 with

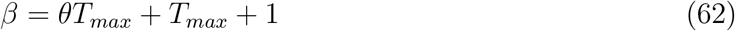

and the initial condition *p_θ_*. Thus, for any *θ* ∈ (0, 1), there exist a time *T_θ_ >* 0, defined as in (47) with *K_θ_* instead of *K*, and a unique solution *p*̃ ∈ 𝒞 ^1^([0*, T_θ_*]*, L^∞^*(ℝ) ∩ *L*^1^(ℝ)) of (3) with initial condition *p_θ_*, such that

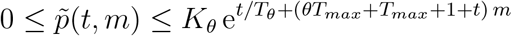

for all *t* ∈ [0*, T_θ_*] and a.e. *m* ∈ ℝ_−_, and supp *p*̃(*t, ·*) ⊂ ℝ_−_ for every *t* ∈ [0*, T_θ_*]. Therefore, for any *θ* ∈ (0, 1), problem (3) with initial condition *p*_0_ has a solution *p* ∈ 𝒞^1^([0*, θ T_max_* + *T_θ_*]*, L^∞^*(ℝ) *∩ L*^1^(ℝ)) such that, for all *t* ∈ [*θ T_max_, θ T_max_* + min(*T_θ_*, 1)]

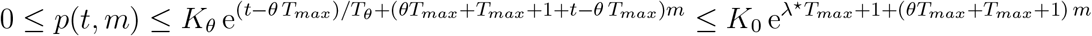

for a.e. *m* ∈ ℝ_−_, and supp *p*(*t*,) ⊂ℝ_−_.

On the other hand, from (61) and from the definitions (62) of *β* and (47) of *T_θ_ >* 0 with *K_θ_* instead of *K*, it follows that

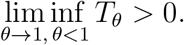

Therefore, there exist *θ′* ∈ (0, 1) and *T′* ∈ (*T_max_, T_max_* + 1) for which problem (3) with initial condition *p*_0_ has a solution *p* ∈ 𝒞^1^([0*, T′*]*, L^∞^*(ℝ) *∩ L*^1^(ℝ)) such that, for all *t* ∈ [*θ*′*T_max_, T′*],

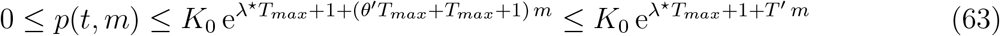

for a.e. *m* ∈ ℝ_−_, together with supp *p*(*t, ·*) ⊂ ℝ_−_. Furthermore, (60) (remember that *β*_0_ = *T_max_* + 1) implies that, for all *t* ∈ [0*, θ′T_max_*] and a.e. *m* ∈ ℝ_−_,

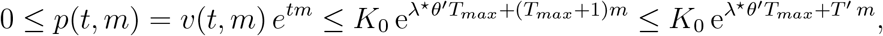

and supp *p*(*t*,) ⊂ ℝ_−_ for all *t* [0*, θ′T_max_*]. Together with (63) in [*θ′T_max_, T′*] ℝ_−_, it follows that the solution *p* satisfies (57) for all *T* ∈ (0*, T′*]. Finally, one infers that *m*̅ ∈ 𝒞 ([0*, T′*)). The fact that *T′* is larger than *T_max_* contradicts the definition of *T_max_*.

As a conclusion, *T_max_* = +*∞* and, from (57) holding for any *T >* 0, property (20) holds with Γ_*α,T*_ = *C*_max(*α,T*)_. From the equation (3) itself and from (10), it also follows that *|∂_t_p*(*t, m*)*|* decays faster than any exponential function as *m → −∞* in the sense that (20) holds for *|∂_t_p*(*t, m*)*|* as well. The proof of Theorem 3.3 is thereby complete.

### 6.3 Proof of the results on the stationary states of (24) and (33)

#### Proof of Proposition 3.5

Let us first show that sup{supp *p_∞_*} = 0. Assume not. Then there is *δ* > 0 such that

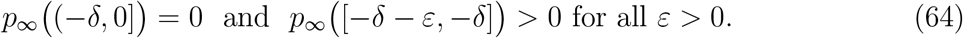

Consider now any nonnegative continuous function *ϕ*: ℝ → ℝ with compact support included in [−*δ*, 0], and such that

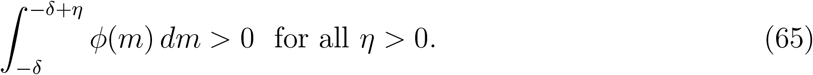

Then, using (10) with *J̄ L*^1^(∈), and the fact that the function *y* ⟼ *∫*_ℝ_ *J_y_* (*m* – *y*) *ϕ*(*m*) *dm* is continuous (and bounded), it follows (as in the derivation of (33)) that

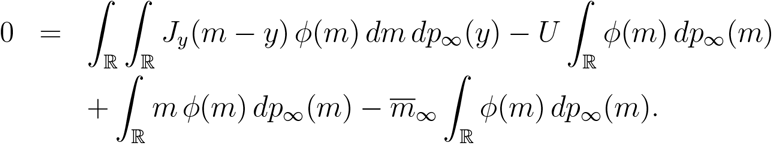

Since the measure *p*_∞_ is supported in (−*∞*, −*δ*] and the continuous function *ϕ* is supported in [−*δ*, 0], one infers from the previous formula that

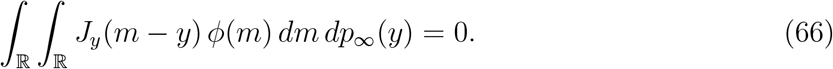

From the assumption (18) on the positivity and continuity of *y* ⟼ *S_y_* on (*−∞*, 0), there is *ε* ∈ (0, *δ*) such that 2*ε* < *S_y_* for all *y* ∈ [−*δ* − *ε*, −*δ*]. It then follows from (66) and the nonnegativity of *J_y_*, *ϕ* and *p*_∞_ that

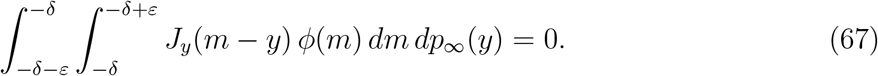

On the other hand, for each *y* ∈ [−*δ* −*ε*, −*δ*] and each *m* ∈ [−*δ*, −*δ* + *ε*], one has 0 ≤ *m* − *y ≤* 2*ε < S_y_*. Hence, for every *y* ∈ [−*δ −ε*, −*δ*], one has *J_y_*(· − *y*) > 0 a.e. in [−*δ*, −*δ* + *ε*]. Together with (65) and the nonnegativity of *ϕ*, it follows that

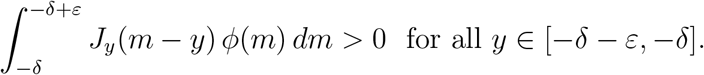

Together with (67) and the nonnegativity of *p*_∞_, one gets that *p_∞_*([–*δ* –*ε*, –*δ*]) = 0, a contradiction with (64). Therefore, one has shown that

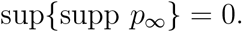

Lemma 4.5 in [17]^2^ then implies that the CGF *C*_∞_ of *p*_∞_ satisfies 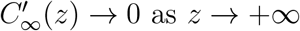.

Finally, let us prove that *m̄_∞_* ≥ −*U*. Using equation (33) satisfied by *C*_∞_ and the fact that 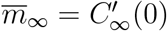, we obtain that

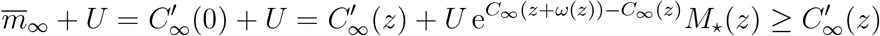

for all *z* ≥ 0. Thus, by passing to the limit as *z →*+ ∞, one immediately gets that *m̄*_∞_+*U* ≥ 0. The proof of Proposition 3.5 is thereby complete.

#### Proof of Proposition 3.6

Equation (33) satisfied by *C*_∞_ can be rewritten as

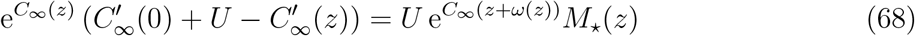

for all *z* ≥ 0. On the one hand we have

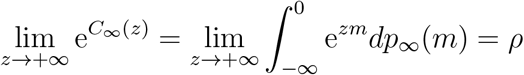

from the definition of *ρ* ∈ [0, 1] in (27). On the other hand, we know from the previous proposition that 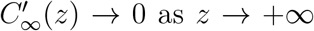. By passing the limit as *z* → +∞ in (68), we get that the right-hand side has a limit and that

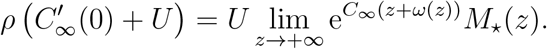

Since

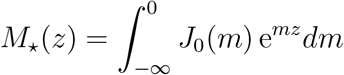

is the MGF of the mutation kernel *J*_0_ at the optimal fitness, with *J*_0_ ∈ *L*^1^(ℝ), one has lim_*z*→+*∞*_ *M*_*⋆*_(*z*) = 0. Moreover, the function 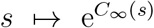 is continuous in [0, +*∞*) and converges to *ρ* ∈ [0, 1] at +∞. Thus, it is bounded in [0, +*∞*). Therefore, we get that 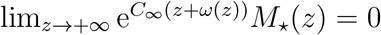 and

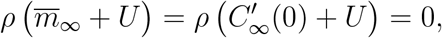

that is to say that *ρ* = 0 or *m̄_∞_* = −*U*.

#### Proof of Proposition 3.7

Let us consider the MGF of *p*_∞_, namely the function defined in ℝ_+_ by

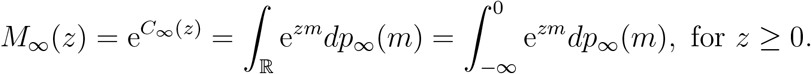

Equations (33) and (68) with 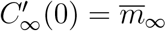 can be rewritten as

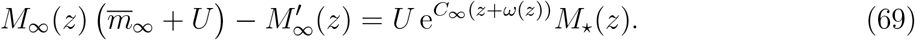

We then consider separately the cases 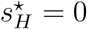 and 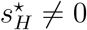, where 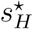 is defined in (28).

*First case:* 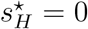. This case means that

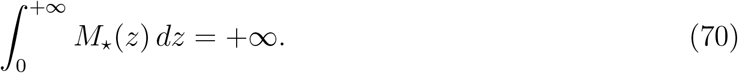

Meanwhile, due to (8), (9) and (11), the function *ω* necessarily satisfies some properties, whose proof is postponed in Appendix B below:

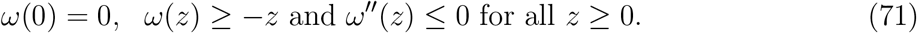

Thus, it follows the function *z* ⟼ *ω*(*z*) + *z* is nonnegative and nondecreasing. It then has a limit *l* ∈ [0, +*∞*] as *z* → +∞. Assume first here that

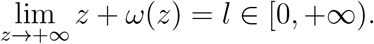

Thus, 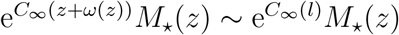 as *z* → +∞ and

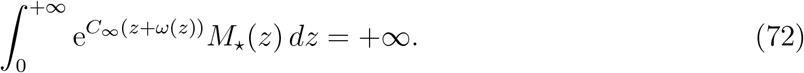

Since 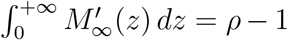 and *M*_∞_ (*z*) > 0 for all *z* ∈ [0, +*∞*), it follows from (69) and (72) that *m̄*_∞_ + *U* > 0. From Proposition 3.6, this means that *ρ* = 0.

Assume now that

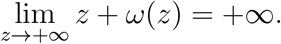

Since *C*_∞_ is nonincreasing in [0, + ∞) from its definition (32) and since *ω* is nonpositive in [0, +∞) by (12) and (71), we get that 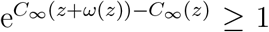 for all *z* ≥ 0. Thus, using (70), one infers that

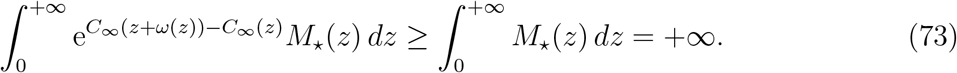

Let us then show that *ρ* = 0. If *m̄_∞_* + *U* > 0, then Proposition 3.6 yields *ρ* = 0. Assume now that *m̄_∞_* + *U* = 0, in other words 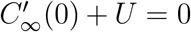. By integrating (33) over (0*, A*) with *A >* 0, we obtain that

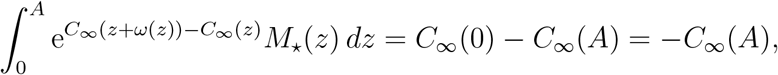

that is,

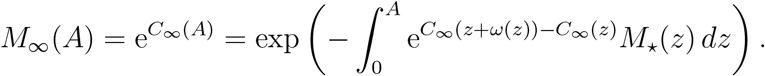

Knowing that lim_*A*→+∞_ *M*_∞_(*A*) = *ρ* and using (73), we finally get that

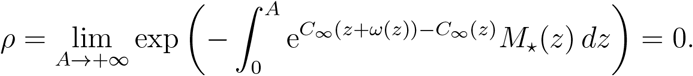

*Second case* 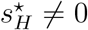. This case means that

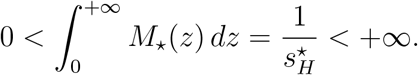

Since *C_∞_* is nonpositive by (32) and *p_∞_*((−*∞*, 0]) = 1, we also have that

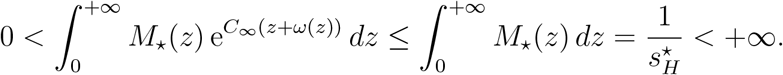

Using the inequality *m̄*_∞_ + *U* ≥ 0 and integrating (69) over (0, +*∞*) yields

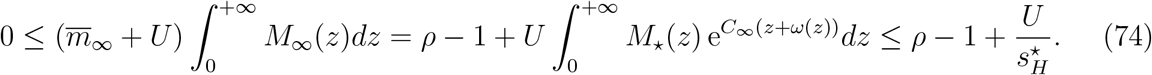

Therefore, if 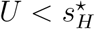, then *ρ* > 0, and *m̄*_∞_ = –*U* by Proposition 3.6.

Let us now consider the set

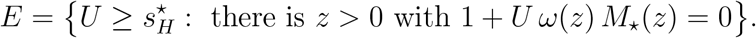

Remember that *M*_⋆_(*z*) > 0 for all *z* ≥ 0. Furthermore, as *ω* satisfies (12) and (71), there is *z*_0_ > 0 such that *ω* < 0 in [*z*_0_, +∞). Therefore, for every 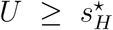 such that *U* > −1/(*ω*(*z*_0_)*M*_⋆_(*z*_0_)) (> 0), there holds 1 + *U ω*(*z*_0_) *M*_⋆_(*z*_0_) < 0 and, by continuity of *ω* and *M*_⋆_ and since *ω*(0) = 0, there is *z*_1_ > 0 such that 1 + *U ω*(*z*_1_) *M*_⋆_(*z*_1_) = 0. As a consequence, the set *E* is not empty and

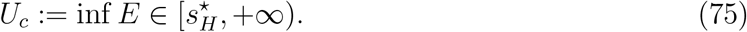

Pick any *U* > *U_c_*. We want to show that *ρ* = 0 and *m̄_∞_* > −*U*. Assume by contradiction that *m̄*_∞_ = −*U*, that is, 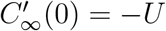. Using (33), we obtain that, for all *z* ≥ 0,

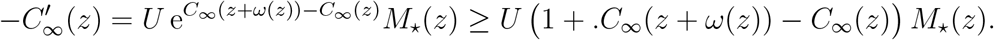

Using (71) and the convexity of *C_∞_* in [0, +*∞*), one infers) that, for all *z* ≥ 0, 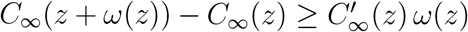 hence 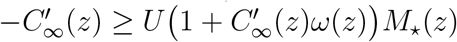, that is,

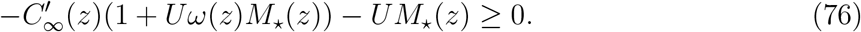

On the other hand, as *U > U_c_*, there are 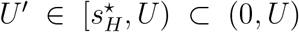 and *z′ >* 0 such that 1 + *U′ω*(*z′*) *M*_⋆_(*z′*) = 0, hence *ω*(*z′*)*M*_⋆_(*z^t^*) = 1*/U′ <* 0 and 1 + *U ω*(*z′*) *M*_⋆_(*z′*) *<* 0. Again by continuity, one infers the existence of *z*_1_ *>* 0 such that

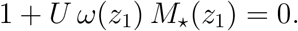

Formula (76) at *z* = *z*_1_ yields –*U M*_⋆_(*z*_1_) 0, which is ruled out since *U* > 0 and *M_⋆_*(*z*_1_) > 0. Thus, *m̄*_∞_ > *–U*, and *ρ* = 0 by Proposition 3.6. The proof of Proposition 3.7 is thereby complete.

#### Proof of Corollary 3.8

Using (31), it is easily seen that 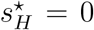 is equivalent to *n* ≤ 2. Furthermore, in the Gaussian Fisher’s geometric model, *ω*(*z*) = −*λz*^2^/(1 + *λz*). Thus,

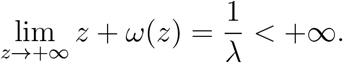

Together with Proposition 3.7, we get the conclusion (i) of Corollary 3.8 in case *n* ≤ 2.

In case *n* > 2, then 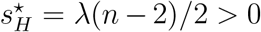. Using again Proposition 3.7, we immediately get that *ρ* > 0 and *m̄*_∞_ = –*U* if *U < λ*(*n* – 2)/2.

Let us show that the same result holds good with the large inequality 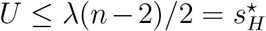 (still with *n* > 2). First of all, by using Proposition 3.6 and equation (69) multiplied by *ρ*, one has

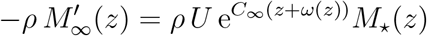

for all *z ≥* 0. Hence, afther integration over (0, +*∞*), one infers that

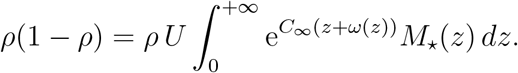

Therefore, *ρ <* 1. As a consequence, the function *C*_∞_ defined in (32) (with *p*_∞_ written as in (27)) is (strictly) decreasing in [0, + ∞) and thus negative in (0, + ∞). Notice that these properties hold under the general assumptions of Proposition 3.7. On the other hand, since, here in the Gaussian Fisher’s geometric model, *ω*(*z*) = − *λz*^2^/(1 + *λz*) for all *z* ≥ 0, one has *z* + *ω*(*z*) > 0 for all *z* > 0, hence e^*C∞*(*z*+*ω*(*z*))^ < 1 for all *z* > 0. Together with the positivity of *M_*_*(*z*), it follows that

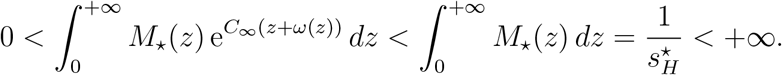

As a consequence, if 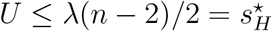, then, as in (74), one gets that

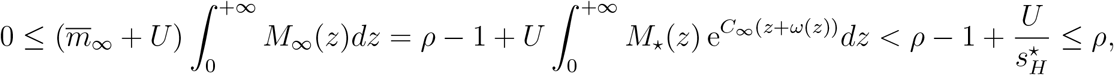

hence *ρ* > 0, and *m̄*_*∞*_ = –*U* by Proposition 3.6.

Lastly, still in case *n* > 2, a straightforward computation, using (13), implies that the quantity *U_c_* defined in (75) is equal to

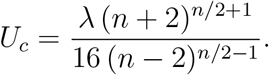

Conclusion (ii)-(b) of Corollary 3.8 then follows immediately from Proposition 3.7.

## 7 Proof of technical lemmas

This section is devoted to the proof of the technical lemmas, namely Lemmas 6.1, 6.4 and 6.5, used in the proof of the main theorems in the previous sections.

### 7.1 Comparison principle

#### Proof of Lemma 6.1

Let *τ*, *w*, *h*_1_ and *h*_2_ satisfy the assumptions of the lemma. With a slight abuse of notations, we write 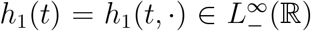 and 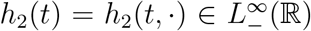 for *t* ∈ [0, *τ*]. Set

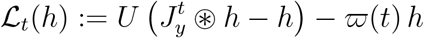

for *t* ∈ [0, *τ*] and 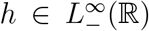. For any such 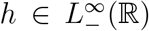, one has supp ℒ_*t*_(*h*) ⊂ ℝ– and, from (8), (10), (37) and similar calculations as in (52), there holds

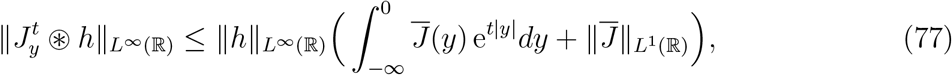

hence, 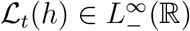. Then let us denote

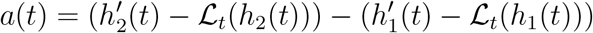

for *t* ∈ [0, *τ*]. Notice that supp *a*(*t*) ⊂ ℝ_−_ for every *t* ∈ [0, *τ*]. Furthermore, for any *i* ∈ {1, 2}, any *t* ∈ [0, *τ*] any sequence (*t_n_*)_*n*∈ℕ_ in [0, *τ*] converging to *t*, any *m* ∈ ℝ_−_ and any *n* ∈ ℕ, one has

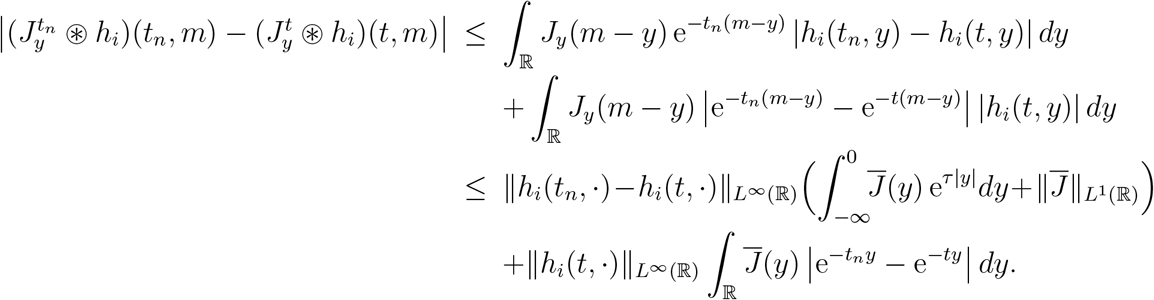

Therefore, 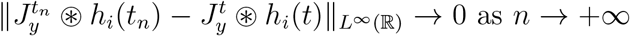 as *n* → + ∞ from the assumptions on *h_i_* and *J̄* and from Lebesgue’s dominated convergence theorem. Finally, one infers that the maps *t* ↦ *ℒ_t_*(*h_i_*(*t*)) belong to 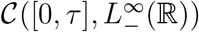, and that 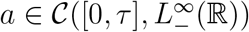.

Now define, for *t* ∈ [0, *τ*],

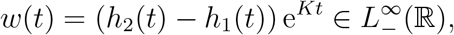

with

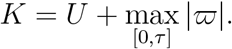

It is straightforward to check that *w* is a solution of the ordinary differential equation

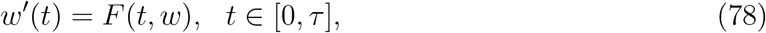

in 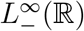, for some function 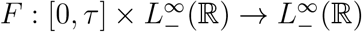 defined by

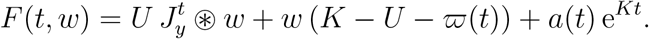

As above, the function *F* is continuous in 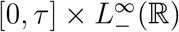. Furthermore, for any *t* ∈ [0, *τ*] and *w*, 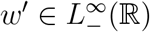, one has, as in (77),

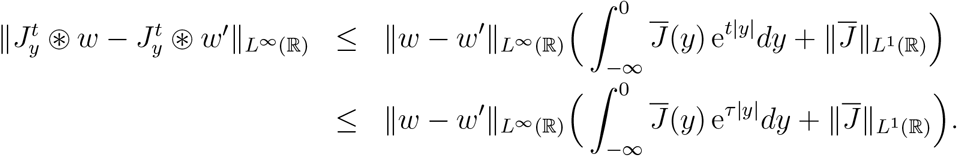

As a consequence, the function *F* is Lipschitz-continuous with respect to *w* uniformly in *t* ∈ [0, *τ*]. We can then define 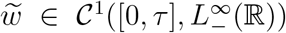 as the unique solution of 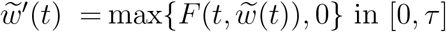 with *w̃*(0) = *w*(0), that is,

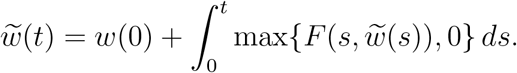

We have *w̃*(*t*) ≥ *w*(0) and *w*(0) ≥ 0 a.e. in ℝ by assumption. Additionally, from (42), there holds *a*(*t*) ≥ 0 a.e. in ℝ_–_, for all *t* ∈ [0, *τ*]. As a consequence, and since *K* – *U* – ϖ (*t*) ≥ 0 for all [0, *τ*], one infers that, for all *t* ∈ [0, *τ*], *F* (*t*, *w̃*(*t*)) ≥ 0 a.e. in ℝ_−_. We deduce that *w̃* is also a solution of the equation (78) satisfied by *w*. From the Cauchy-Lipschitz theorem, we deduce that, for all *t* ∈ [0, *τ*], *w*(*t*) = *w̃*(*t*) ≥ 0 and therefore *h*_1_(*t*,·) ≤ *h*_2_(*t*,·) a.e. in ℝ. The proof of Lemma 6.1 is thereby complete.

### 7.2 Existence for the Cauchy problem

#### Proof of Lemma 6.4

Let *υ* ∈ *E*. We know by definition of the set *E* and Lebesgue’s dominated convergence theorem that the function

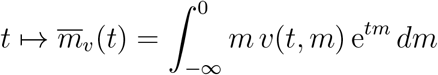

is continuous in [0, *T*]. Problem (50) can then be written as an ordinary differential equation

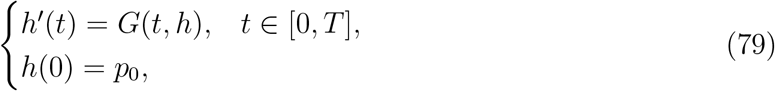

with

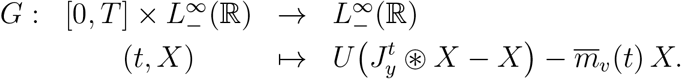

The function space 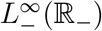 is a Banach space for the uniform norm ‖ ‖_∞_ and as in the proof of Lemma 6.1 above, the function *G* is continuous in 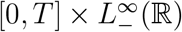 and Lipschitz continuous with respect to *X* uniformly in *t* ∈ [0, *τ*]. Therefore, the Cauchy-Lipschitz theorem yields the existence and uniqueness of a solution 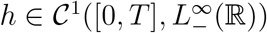 of problem (79).

### 7.3 Mass preservation and estimates on mean fitness

#### Proof of Lemma 6.5

Let us first show (58). We consider

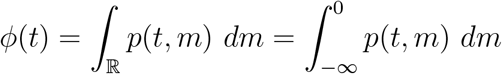

for *t* ∈ [0, *T_max_*). This quantity is a well-defined nonnegative real number due to (57) and the definition of *T_max_*. By using (57) for every *T* ∈ [0, *T_max_*), integrating (3) against *m* over ℝ_−_, we obtain that *ϕ* is of class *𝒞*^1^([0, *T_max_*)) and, for every *t* ∈ [0, *T_max_*),

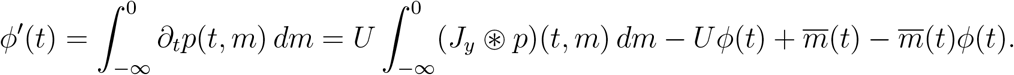

From assumption (8), we have

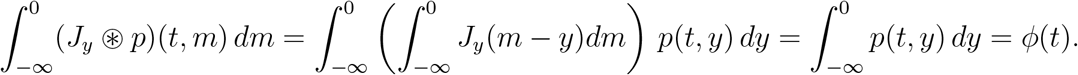

Finally, *ϕ′*(*t*) = *m̄*(*t*)(1 − *ϕ*(*t*)) for all *t* ∈ [0, *T_max_*). From assumption (5), there holds *ϕ*(0) = 1, and since *m̄* ∈ *𝒞*([0, *T_max_*)), it follows immediately that *ϕ*(*t*) = 1 for all *t* ∈ [0, *T_max_*).

Let us now turn to the proof of (59). By integrating (3) between 0 and *t* ∈ [0, *T_max_*), multiplying by *m*, integrating over ℝ_−_ and using (57) and Fubini’s theorem, we get that, for every *t* ∈ [0, *T_max_*),

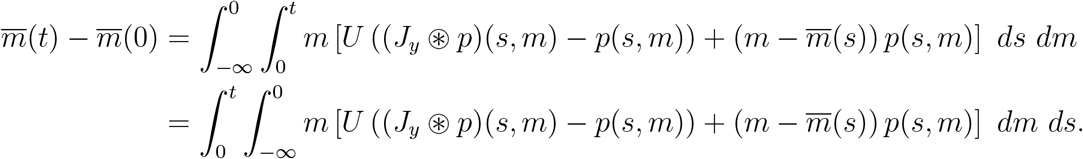

From (8), (10), (57), (58) and Fubini’s theorem, one infers that, for every *s* ∈ [0, *t*] (⊂[0, *T_max_*)),

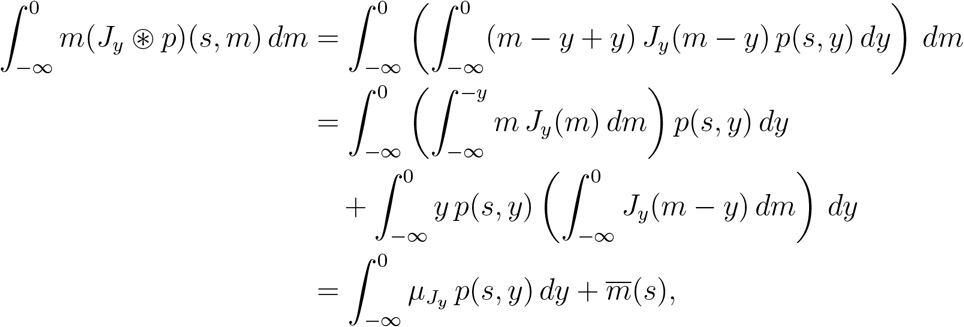

with

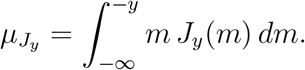

Then, using the fact that the functions 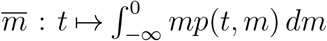 and 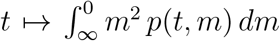 are continuous in [0, *T_max_*), we deduce that, for every *t* ∈ [0, *T_max_*),

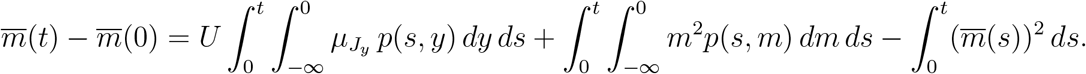

Then, by the Cauchy-Schwarz inequality and (58), we have

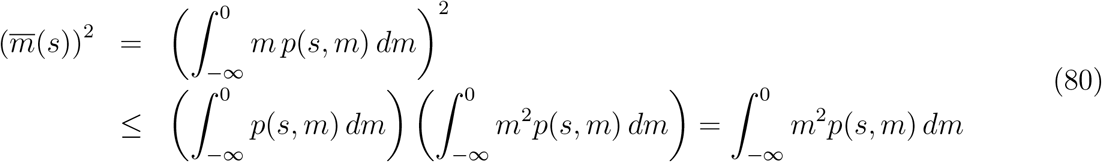

for all *s* ∈ [0, *t*] (⊂[0, *T_max_*)). Therefore,

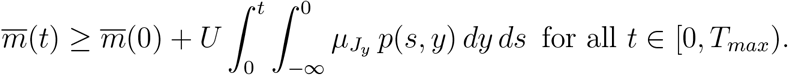

Finally, using assumptions (8) and (10), we have, for all *y* ≤ 0,

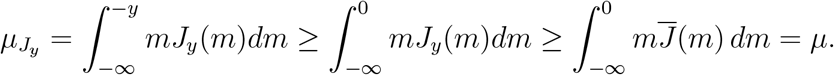

Thus, since *p* is nonnegative and 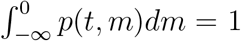 for all *t* ∈ [0, *T max*), we get that *m̄*(*t*) ≥ *m*(0) + *t U μ* for all *t* ∈ [0, *T_max_*). The proof of Lemma 6.5 is thereby complete.

## Appendix A: Interpreting condition (29) on 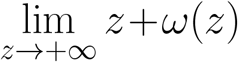

We here show that the assumption (29) of Proposition 3.7, namely lim_*z*→+*∞*_(*z* + *ω*(*z*)) ∈ [0, +*∞*), means that any parent can give mutant offspring with the optimal fitness 0, that is, sup{supp *J_y_*} = –*y* for every *y* ≤ 0. To do so, differentiating equation (15) with respect to *z*, we have (as in (17)):

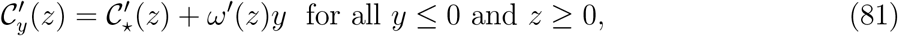

where the *𝒞*^∞^(ℝ_+_) functions *𝒞_y_* and *𝒞_⋆_* are defined by (16). Now, [17, Lemma 4.5] yields 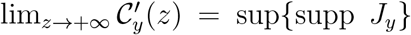 and 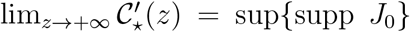. Passing to the limit *z* → +*∞* in (81), it follows that lim_*z*→+*∞*_ ω′(*z*) exists and

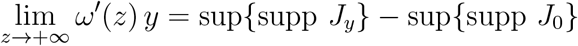

for all *y* ≤ 0. Assumption (29) then implies that *ω′*(*z*) → −1 as *z* → +*∞*, hence

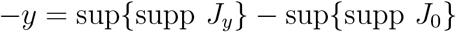
for all *y* ≤ 0. On the other hand, assumption (9) yields in particular sup{supp *J*_0_} ≤ 0. If sup{supp *J*_0_} = −*α* < 0, then sup{supp *J_y_*} = −*y* − *α* for all *y* ≤ 0, contradicting assumption (18) for *y* > 0 small. Therefore, sup{supp *J*_0_} = 0, and

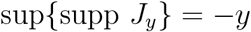

for all *y* ≤ 0. In other words, any parent can give mutant offspring with the optimal fitness 0.

## Appendix B: Proof of property (71) on *ω*

First of all, by taking *z* = 0 in (11), we immediately get that

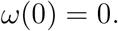

Second, since

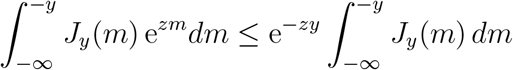
for all *y ≤* 0 and *z ≥* 0, it follows from (8) and (9) that

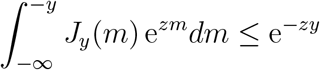

for all *y* ≤ 0 and *z* ≥ 0. Then, using (11), we get that *M* _⋆_ (*z*) e^*ω*(*z*)*y*^ ≤ e^*−zy*^ and, for every *y* < 0, we obtain that (ln *M*_⋆_(*z*))*/y* + *ω*(*z*) ≥ − *z* for all *z* ≥ 0. The limit as *y* → –*∞* then yields *ω* (*z*) ≥ − *z* for all *z* ≥ 0.

Third, observe that assumption (11) yields

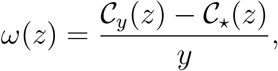

for all *y* < 0 and *z* ≥ 0. Thus, as *𝒞_y_* and *𝒞*_⋆_ belong to *𝒞^∞^*(ℝ_+_), the function *ω* belongs to *𝒞^∞^*(ℝ _+_) too. In order to show the concavity of *ω*, it is then sufficient to prove that *ω*″ ≤ 0 in ℝ_+_. But since *𝒞_y_* is convex in ℝ_+_ for every *y* ≤ 0, the previous displayed formula implies that

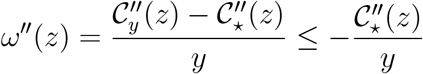
for all *y* < 0 and *z* ≥ 0. The limit as *y* → –*∞* yields *ω″*(*z*) ≤ 0 for all *z* ≥ 0, and the proof of (71) is thereby complete.

## Footnotes

1 With a slight abuse of notation, we also use this notation for functions which are not necessarily in *E*. Notice that the set *E* is complete for the topology induced by *‖ ‖_E_*.

2 It is immediate to see that the proof and the conclusion of [17, Lemma 4.5] hold good even if the nonnegative probability measure *p_∞_* is not in *L^∞^*(ℝ).

